# *Stemona* genomes illuminate fatty acid partitioning between seeds and elaiosomes mediating wasp dispersal

**DOI:** 10.64898/2026.02.19.706862

**Authors:** Tianquan Yang, Nathanael Walker-Hale, Fengmao Yang, Chunxia Zeng, Zhenshan He, Guillaume Chomicki, Wei Xu, Gao Chen

## Abstract

- Seed dispersal by wasps (vespicochory) is documented in five angiosperm families, with phylogenies suggesting that most vespicochorous lineages evolved from ant-dispersed (myrmecochorous) ancestors. While recent work has identified cues attracting wasp dispersers, the molecular basis remains unclear.
- To shed light on the molecular basis of vespicochory, we generated chromosome-level genomes for the wasp-dispersed *Stemona tuberosa*, and its close ant-dispersed relative *S. mairei*. Combining comparative genomic, transcriptomic, lipidomic, and functional analyses, we ask (i) how chemical elaiosome differentiation occurs during seed development, and (ii) how key mutualism-associated genes evolved in *Stemona*.
- We show that elaiosomes up-regulate stearoyl-ACP Δ9 desaturases (SAD) and accumulate oleic acid and 1,2-diolein, which serve both as food rewards and as precursors for CER1/3-mediated biosynthesis of (Z)-9-tricosene, the key wasp attractant. By contrast, high expression of FatB and DGAT1 in seeds is associated with medium-chain fatty acids (MCFAs) production. Comparative genomics indicates conservation of these genes across *Stemona*, with *S. tuberosa*-specific FatB expansion and SAD/CER1/3 repertoire divergence.
- Our work supports a model for elaiosome-seed fatty-acid differentiation linking recruitment and nourishment in vespicochory. Given that oleic acid and 1,2-diolein are also key cues in myrmecochory, this shared biochemical basis may help explain how vespicochory evolves from ant-dispersed ancestors.

## Introduction

Seed dispersal is a critical process to community structure and stability, and ultimately biodiversity maintenance (Janzen, 1970; Howe and Smallwood, 1982; Willson and Traveset, 2000). Most seed plants are dispersed by animals, involving diverse strategies to attract, reward, or attach to animal dispersers (Tiffney, 2004; Jordano et al., 2007; Schupp *et al*., 2010; Farwig and Berens, 2012), but seed dispersed by invertebrates has received comparatively less attention, with the exception of seed dispersal by ants (myrmecochory). Seed dispersal by invertebrates includes rare endozoochorous dispersers by such as giant flightless grasshoppers (wetas, Duthie *et al*., 2006), cockroaches (Uehara & Sugiura, 2017), camel crickets (Suetsugu, 2018) and red slugs (Türke *et al*., 2012). Other invertebrate seed dispersal involves non-rewarding seeds carried by dung beetles (Midgley et al., 2015), or seeds with elaiosome rewards dispersed by flies (Yu *et al*., 2025) or wasps (Chen *et al*., 2018a).

Dispersal by ants, flies and wasps rely on lipid-rich seed appendages called elaiosomes that serves to recruit and reward the dispersers (Mark & Olesen, 1996; Lengyel *et al*., 2010; Chen *et al*., 2018a; Yu *et al*., 2025). Elaiosomes have evolved independently in ∼11,000 plant species from over 80 angiosperm families (Edwards *et al*., 2006; Lengyel *et al*., 2010). Ants and wasps are attracted by the elaiosome-produced signal molecules and quickly remove the seeds (Karnish, 2024; Chen *et al*., 2018). These cues are lipids, especially oleic acid and diglyceride 1,2-diolein (a diacylglycerol with oleic acid at positions 1 and 2) are key signalling compounds to trigger corpse-carrying behaviour (Skidmore & Heithaus, 1988; Brew *et al*., 1989; Fischer *et al*., 2008; Pfeiffer *et al*., 2010; Boieiroa *et al*., 2012). These compounds are byproducts of ant decomposition and are part of the conserved necrophoric pathway through which ants identify and remove dead nestmates to prevent disease spread. In addition to recruiting ants, the high oleic acid content in elaiosomes may provide nutrients particularly important for these seed dispersers as rewards (Lanza *et al*., 1992; Hughes *et al*., 1994; Fischer *et al*., 2008). Wasp-addressed elaiosomes are typically larger and use distinct compounds for recruitment, notably lipid-derived cuticular hydrocarbons such as (Z)-9-tricosene in *Stemona tuberosa* (Stemonaceae, Pandanales, Monocotyledons, 2*n* = 14) (Chen *et al*., 2018) and short-carbon-chain volatiles mimicking herbivore damage as in *Aquilaria sinensis* (Thymelaeaceae) (Qin *et al*., 2022).

Beyond attraction compounds, little is known on the molecular mechanism underlying vespicochory. Using phylogenies, chromosome-level genome sequencing for wasp and ant-dispersed *Stemona* species, transcriptomic, lipidomic, and functional analyses we ask the following questions: (i) how chemical elaiosome differentiation occurs during seed development, and (ii) how the underlying genetic toolkit has evolved in *Stemona*. We identify key genes involved in the lipid differentiation of the elaiosome from the seed, and show how their differential expression leads to the accumulation of attracting and rewarding compounds for insect dispersers. Comparative genomics reveals a shared genetic toolkit for wasp and ant dispersal, showing that only a small step is required to shift from myrmecochory to vespicochory.

## Materials and Methods

### Genome sequencing and assembly

*Stemona tuberosa* was collected from cultivated plants at Kunming Botanical Garden, Kunming Institute of Botany, Yunnan Province, China. *S. mairei* was collected from wild plants in Daju County, Yunnan Province, China. Young and healthy leaves of both species were collected and immediately flash-frozen in liquid nitrogen. Genome size was estimated by flow cytometry (Doležel et al., 2007) on fresh leaf tissue from the same individuals. Subsequently, high-coverage resequencing was performed for both species, to estimate genome size. For *S. tuberosa*, genomic DNA was isolated using a Qiagen DNA purification kit (Qiagen, Darmstadt, Germany), quality assessed via NanoDrop (Thermo Fisher Scientific, Wilmington, DE, USA) and concentration with a Qubit fluorometer (Thermo Fisher Scientific). DNA libraries were constructed following the manufacturer’s protocol (Illumina), and sequenced on an Illumina HiSeq 4000 platform with 150 bp paired-end. For *S. mairei*, total genomic DNA was isolated using the DNA Marker D15000; MD110-02 (Tiangen, Beijing, China) and quality-checked as above. Libraries were prepared using the OnePot Pro DNA Library Prep Kit V4 (Yeasen Biotechnology, Shanghai, China), and sequenced was on a DNBSEQ-T7 platform with 150 bp paired-end reads. Raw reads were quality-filtered with fastp (Chen *et al*., 2018b), and genome size and complexity were estimated by counting k-mers (k = 19) using KMC v.3.2.2 (Kokot *et al*., 2017) and fitting the resulting spectra with Genomescope v.2.0 (Ranallo-Benavidez *et al*., 2020), assuming diploidy for both species (Fig. S1).

Long read sequencing was performed for both species, and its integrity was assessed by pulsed-field capillary electrophoresis (Agilent Technologies, Santa Clara, CA, USA). Approximately 8 µg of DNA was sheared using a Megaruptor 3 system (Diagenode SA., Seraing, Belgium), and the fragments were concentrated with AMPure PB beads (Pacific Biosciences, Menlo Park, CA, USA). A SMRTbell sequencing library with ∼15–20 kb inserts was constructed from the high-quality DNA and sequenced on the PacBio Sequel II platform in Circular Consensus Sequencing (CCS) mode. HiFi reads were generated from the raw subreads using the CCS workflow (v.6.3.0) under default parameters (Wenger *et al*., 2019).

To generate a chromosome-scale assembly of both species, we carried out Hi-C sequencing. Young and healthy leaves were cross-linked with formaldehyde at room temperature and genomic DNA isolated from the nuclei was digested with HindIII restriction enzyme. Fragments with sticky ends were then end-repaired and marked with biotinylated nucleotides, and then adjacent blunt ends were ligated to form a circular molecule. Subsequently, these circular molecules were purified and sheared to 300-500 bp, and sequenced on an Illumina HiSeq 4000 platform. For chromosome-scale genome scaffolding, raw Hi-C data were processed into a contact matrix using Juicer v.1.6 (Durand *et al*., 2016) and contigs were then scaffolded with 3D-DNA v.180114 (Dudchenko *et al*., 2017), and manually curated in Juicebox v1.9.8 (Durand et al., 2016; Fig. S2). Chromosome-scale assemblies were polished with TGS-GapCloser v.1.2.1 (Xu *et al*., 2020) and NextPolish2 (Hu *et al*., 2024).

### Genome annotation and comparative genomics

The detailed methodology for this section is reported in the Online *Supplementary Methods*. In short, repetitive elements were annotated de novo using EDTA v2.2.2 (Ou *et al*., 2019) and softmasked with RepeatMasker v4.18 (Smith *et al*., 2013). Protein-coding genes were annotated by integrating ab initio predictions (SNAP, Korf, 2004; AUGUSTUS v3.5.0, Stanke *et al*., 2006), RNA-seq transcript assemblies (Trinity v2.15.2, Grabherr *et al*., 2011; HISAT2 v2.2.1 and StringTie v2.1.4, Pertea *et al*., 2016; PASA v2.5.3, Haas *et al*., 2008), and homologous protein evidence using MAKER2 v3.01.03 (Holt & Yandell, 2011) and EVidenceModeler v2.1.0 (Haas *et al*., 2008); full details are in *Supplementary Methods*. Macrosynteny was assessed with MCScan (JCVI; Tang *et al*., 2024) and visualised with circos v0.69-10 (Krzywinski *et al*., 2009). A phylogenomic dataset for Pandanales was assembled from annotated genomes and published transcriptomes (Tables S1-S2) following the pipeline of Yang & Smith (2014), with gene trees inferred using IQ-TREE v3.0.1 (Wong *et al*., 2025), a species tree estimated with ASTRAL-IV v1.23.4.6 (Zhang *et al*., 2025), and divergence times inferred with mcmctree in PAML v4.10.9 (Yang & Rannala, 2006) using secondary calibrations from Ramirez-Barahona *et al*. (2020) (Supplementary Methods). Gene family evolution was modelled with CAFE5 v1.1 (Mendes *et al*., 2020). Diversifying selection on candidate fatty acid synthesis genes was tested using ABSREL and MEME in HyPhy v2.5.72 (Smith *et al*., 2015; Murrell *et al*., 2012), with foreground branches subtending the MRCA of *Stemona* and *Croomia*, *Stemona*, and *S. tuberosa* (*Supplementary Methods*).

### Transcriptome sequencing and analyses

Transcriptome data from tubers, roots, stems, leaves, flowers, and fruits of *Stemona tuberosa* and *S. mairei* were sampled for genome annotation. To dissect the molecular basis underlying the distinct composition of fatty acid between elaiosomes and seed in *S. tuberosa*, we collected the seeds and corresponding elaiosomes at four developing stages (E1-E4 and S1-S4, see Fig. S3), as single samples.

Total RNA was extracted and purified for each sample using TRIzol (Invitrogen), following the manufacturer’s protocols. The high-quality RNA was sheared into 500 bp fragments and reversely transcribed into cDNA. The cDNAs were then end-repaired and ligated with the Illumina sequencing adapter and sequenced on the Illumina HiSeq 4000 platform with paired-end 150 bp. After quality filtering with fastp (Chen et al., 2018b), clean reads were mapped to the *S. tuberosa* genome using HISAT2 (Kim et al., 2019). Expression levels from uniquely aligning reads were quantified as FPKM using StringTie (Pertea *et al*., 2016).

### Fatty acid profiles in seed and its corresponding elaiosome

Total lipids were extracted from developing elaiosomes (E2 and E4) and seeds (S2 and S4) (**Fig. S4)** and fatty acid composition determined by Gas Chromatography-Mass Spectrometer (GC-MS). For each sample, tissue was ground and transferred into 10 mL glass tubes containing hexane/isopropanol (3:2, v/v) for total lipid extraction. Fatty acid methyl esters (FAMEs) were prepared by adding 2 mL methanol containing 5% H_2_SO_4_ (v/v) and then heating at 85 ℃ for 90 min. FAMEs were dissolved in 1 mL of hexane and analysed on Agilent 7890 gas chromatograph/5975 mass selective detector equipped with a 30 m DB-5MS capillary column. Helium was used as the carrier gas at a constant flow rate of 1 mL/min. The oven conditions included an initial temperature of 40℃ and an initial time of 1 min, 10 ℃/min to 130 ℃, 8 ℃/min to 250℃ for a 10 min bakeout. The inlet temperature was kept constant at 250 ℃, and the MS transfer line was set at 290 ℃. Spectra were acquired by scanning m/z 50-600 in the electron impact (EI) mode for routine analysis. To measure lipid compositions (e.g. DAG, TAG) in elaiosome, we performed UPLC-QTOF/MS-based lipidomic analysis, following (Zhang *et al*., 2022).

### Construction of yeast BY4741 mutant strain *Δole1*

To characterize the function of *S. tuberosa SAD* genes, we constructed the yeast BY4741 (*Saccharomyces cerevisiae*) mutant strain *Δole1* (*MATa ole1Δ::HIS3 leu2Δ0 met15Δ0 ura3Δ0*) by homologous recombination (Fig. S5). Two homologous regions of yeast *OLE3* gene were cloned from genomic DNA of yeast strain BY4741 with specific primers, and an insertion fragment, the HIS3 expression cassette (1364bp), was amplified from the plasmid pFA6a-His3MX6. The three DNA fragments were fused into one fragment by overlap extension PCR. The fused DNA fragment was purified and transformed into yeast strain BY4741 using the protocol of Gietz & Schiestl (2007). The BY4741 *Δole1* could not grow without oleic acid supplementation, and thus *Δole1* strain was grown on the synthetic complete selection medium (-His) with 0.2% oleic acid. All the *Δole1* strain were further validated by PCR with diagnostic primers (Table S3).

### Functional characterization of *S. tuberosa* SAD

Total RNA was extracted from developing seeds and elaiosomes using Trizol (GENEray, SHH, CHN). First-strand cDNA was synthesized using TransScript All-in-One First-Strand cDNA Synthesis kit (TransGen, BJ, CHN). The full-length CDS of two *SAD*s (*Sttub03G0473000.1* and *Sttub07G0192400.1*) was amplified from the cDNA library using Phanta Max Super-Fidelity DNA polymerase (Vazyme, Nanjing, China). The PCR products were cloned into pEASY-Blunt cloning Vector (TransGen, Beijing, China) and verified by Sanger sequencing. After that, the CDS sequence was constructed into the yeast expression vector pYES2.0 using ClonExpress Entry One Step Cloning Kit (Vazyme, Nanjing, China).

Next, the pYES2.0 constructs were transformed into BY4741 *Δole1* using Frozen-EZ Yeast Transformation II Kit (Zymo, USA). Transformants carrying SAD (*Δole1*+*StFAB2* and *Δole1*+*StSAD6*) were first selected on SD-Ura plates supplemented with 0.02% oleic acid, and the transformants carrying empty plasmid were used as the negative control (*Δole1*+empty) (Figs. S4-S5). Positive clones were subsequently grown into liquid SD-Ura medium supplemented with 0.02% sodium oleate for 24 h. Cells were then transferred into the liquid SD-Ura medium supplemented with sodium oleate (0.02%) and 2% galactose to induce transgene expression. After 48 hours, yeast cells were collected for lipid extraction and fatty acid analysis. To assess the effect of oleic acid on MCFAs production, we also collected the *Δole1* cells grown on synthetic complete medium lacking SD-His with 0.2% oleic acids (*Δole1*+ oleic acid).

Yeast cells in stationary phase were collected and lysed in 4 M HCl. After boiling for 10 min, total lipids were extracted with 500 µL hexane/isopropanol (3:2, v/v) and dissolved in 500 µL chloroform. Lipids were then transmethylated with 2 mL of methanol containing 5% H_2_SO_4_ (v/v) at 85 ℃ for 90 min. Finally, FAMEs were analysed by GC-MS as described above. All primers are listed in Table S3.

### Functional characterization of *S. tuberosa* FatB

The full-length CDS of four *FatB* genes (*Sttub01G0002300.1*, *Sttub07G0241600.1*, *Sttub02G0230000.1* and *Sttub02G0229900.1*) was amplified, cloned into pEASY-Blunt cloning Vector (TransGen, Beijing, China) and verified by Sanger sequencing, then subcloned into the yeast expression vector P425 using ClonExpress Entry One Step Cloning Kit (Vazyme, Nanjing, China). P425 constructs were transformed into *S. cerevisiae* strain INVSc1 (*MATa his3Δ1 leu2 trp1-289 ura3-52*). Transformants were selected on SD lacking Leu at 30 ℃, and positive clones grown in liquid SD-Leu at 28 ℃, shaking at 230 rpm. Cells were harvested for lipid analysis as described above.

The same *FatB* gene (*Sttub02G0230000.1*) was additionally transformed into *Arabidopsis thaliana*. Briefly, the full-length CDS of *FatB* was cloned to binary vector pCambia1300 with hygromycin selection gene in plant. Arabidopsis ecotype Col-0 was transformed with *Agrobacterium*-containing *FatB* vector by the floral dip method (Clough & Bent, 1999). Transgenic seedlings of T1 lines were transferred to soil, and T3 seeds were used for fatty acid analysis.

### Substrate preference analysis of *S. tuberosa* DGATs

The full-length CDS of all *S. tuberosa DGAT* genes was cloned and transformed individually into *Saccharomyces cerevisiae* TAG-deficient strain H1246 (*MATα are1-Δ::HIS3 are2-Δ::LEU2 dga1-Δ::KanMX4 Iro1-Δ::TRP1 ADE2 ura3*), for functional complementation using a Yeast Transformation Kit (Coolaber, Beijing). All primers are listed in Table S3. Transformants were selected on SD medium lacking Ura (SD/– Ura) before being grown in liquid SD/–Ura medium (glucose was replaced with galactose) at 30 ℃ with shaking at 200 rpm for 36 h. Cells were harvested by centrifugation for lipid analysis by thin-layer chromatography (TLC). Total lipids (TLs) from yeast were extracted and separated by TLC as described previously (Yang *et al*., 2024). Wild-type INVSc1 and H1246 complemented with the yeast *DGA1* gene served as controls.

To determine the substrate preference of these DGATs, octanoic acid (C8:0), decanoic acid (C10:0), and lauric acid (C12:0) were added to yeast cultures at a concentration of 30 µM. H1246 cells were induced with galactose for 4 days at 30 ℃, and then harvested for TAGs isolation by TLC and fatty acid analysis by GC-MS.

### qRT-PCR analysis

To validate the expression pattern of candidate genes we selected (see Table S4), we used quantitative RT-PCR (qRT-PCR) in different tissues of *S. tuberosa*. Young leaves, developing elaiosomes (E1-E4) and seeds (S1-S4) were collected and immediately flash-frozen in liquid nitrogen. Total RNA was extracted, purified and reversely transcribed into cDNAs as described above. qRT-PCR was performed on a Bio-Rad CFX96 system (CA, USA) using SYBR Green Master Mix (TaKaRa). *S. tuberosa* ACTIN2 (*Sttub01G0312800.1*) gene was used as an internal reference for normalization. The primers used in this study are listed in Table S3.

## Results

### Chromosome-scale assemblies of *Stemona mairei* and *Stemona tuberosa*

In *Stemona*, vespicochory has evolved from myrmecochory at least three times independently, mirroring the general pattern across vespicochorous lineages, with the exception of Thymelaeaceae (Fig. 1a-b). To investigate the molecular basis of vespicochory, we sequenced the vespicochorous *S. tuberosa* and its myrmecochorous close relative *S. mairei*. Flow cytometry estimated the genome size of *Stemona tuberosa* at 0.82 Gb and that of *S. mairei* at 1.27 Gb; k-mer analysis yielded estimates of 899.8 Mb (from 92.5 Gb of short-read data) and 951.7 Mb (from 123.4 Gb), respectively (Fig. S1). A total of 86.7 Gb of HiFi reads and 141.2 Gb of Hi-C reads were generated for *S. tuberosa*, and 31.7 Gb of HiFi reads and 120.5 Gb of Hi-C reads for *S. mairei*.

**Figure 1.**
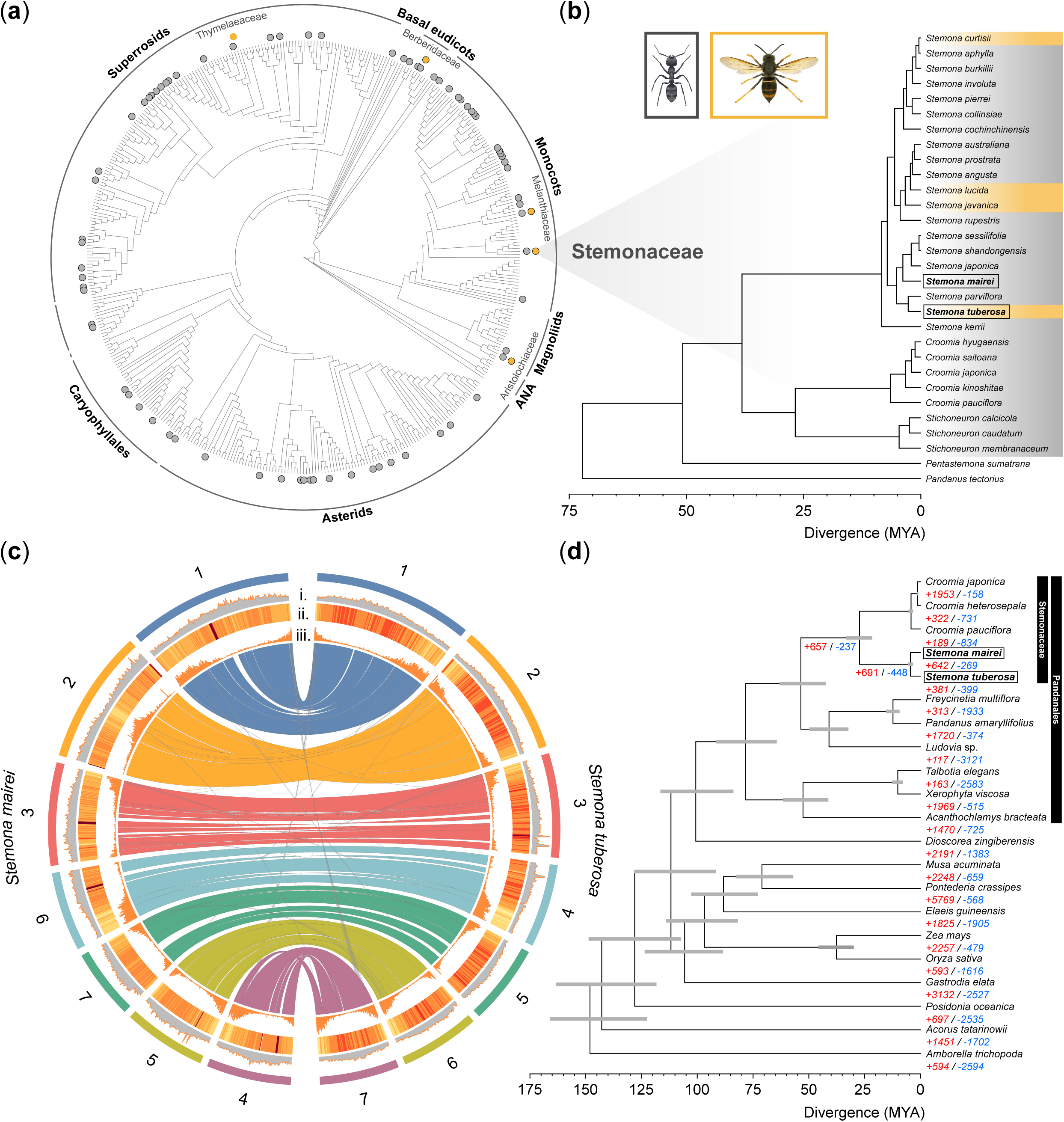
Evolution and comparative genomics of myrmecochory and vespicochory in *Stemona*. (a) the distribution of angiosperm families with reported myrmecochory (grey) and vespicochory (yellow). With the exception of Thymeleaceae, all vespicochory appears to arise from myrmecochorous ancestors. Tree from Ramirez-Barahona et al. (2020), reduced to family-level sampling. (b) evolution of vespicochory in *Stemona*. Dated phylogeny from Chen et al. (2021) showing the distribution of ant-dispersed (grey) and wasp dispersed (yellow) species in *Stemona*. Vespicochory has evolved at least three times from myrmecochorous ancestors. The two species studied in this work are highlighted by boxes. (c) Circos plot comparing chromosome-scale genome assemblies of *Stemona mairei* and *Stemona tuberosa*. i) GC content per 100 Kb ii) repeat count per 100 Kb iii) gene count per 100 Kb. Ribbons show major syntenic blocks between the two genomes. (d) phylogenomic analysis of Pandanales and selected monocots, rooted on *Amborella trichopoda*. Phylogeny with divergence times, grey bars give the 95% Highest Posterior Density (HPD) of the node age. Red (blue) counts give inferred gene family expansions (contractions). *S. tuberosa* and *S. mairei* are highlighted.

The genome assembly of *S. tuberosa* totalled 885.2 Mb with a contig N50 of 84.7 Mb; Hi-C scaffolding anchored 769.4 Mb (86.9%) onto seven pseudochromosomes (Fig. 1c; Fig. S2a). The *S. mairei* assembly totalled 1003.2 Mb (contig N50 of 23.8 Mb), with 937.9 Mb (91.1%) anchored onto seven pseudochromosomes (Fig. 1c; Fig. S2b). BUSCO evaluation yielded completeness scores of 98.4% (1588/1614) for *S. tuberosa* and 98.3% (1585/1614) for *S. mairei*.

Sequence annotation revealed that 66.57% and 74.92% of the genomes consisted of repetitive elements (Table S5), most of which are long terminal repeats (LTR) making up 46.63% and 59.38%, respectively (Table S5). Integrated *ab initio*, homology-based and RNA-seq-aided prediction identified 26,453 protein-coding genes in *S. tuberosa* and 30,875 in *S. mairei* (Table S6), with annotation BUSCO completeness of 98.1% and 97.8%, respectively (Table S7) and ∼77.41% and ∼89.37% protein-coding genes (23,390 and 27,595) were functionally annotated (Table S8).

The two genomes differed by a series of structural rearrangements, including several translocations, nested inversions, and a major inversion of chromosome 2 (Fig. 1c). CAFE5 analysis inferred 381 gene families underwent expansion and 399 underwent contraction in *S. tuberosa* genome, compared with 642 expanding and 269 contracting in *S. mairei*, with the branch subtending the two species sharing a further 691 inferred expansions and 448 inferred contractions (Fig. 1d and Tables S9-S10).

### Fatty acid compositions in elaiosomes and seeds of *S. tuberosa*

Because lipid and their derivatives provide compounds that recruit and reward hymenopteran dispersers (Skidmore *et al*., 1988; Chen *et al*., 2018a), we first characterized the fatty acids compositions in early and mature elaiosomes (E2 and E4) and seeds (S2 and S4) of *S. tuberosa* (see Fig. S3). Elaiosomes were mainly composed of oleic acid (C18:1), making up ∼70% of total fatty acids, followed by palmitic acid (C16:0) and linoleic acid (C18:2) (Fig. 2b). By contrast, seeds accumulated a large proportion of medium-chain fatty acids (MCFAs) including capric acid (C10:0, ∼70% of total lipids), lauric acid (C12:0, ∼17%), and minor amounts of long-chain fatty acids (e.g., C16:0, C18:1 and C18:2, total ∼9.2%) (Fig. 2b).

**Fig. 2.**
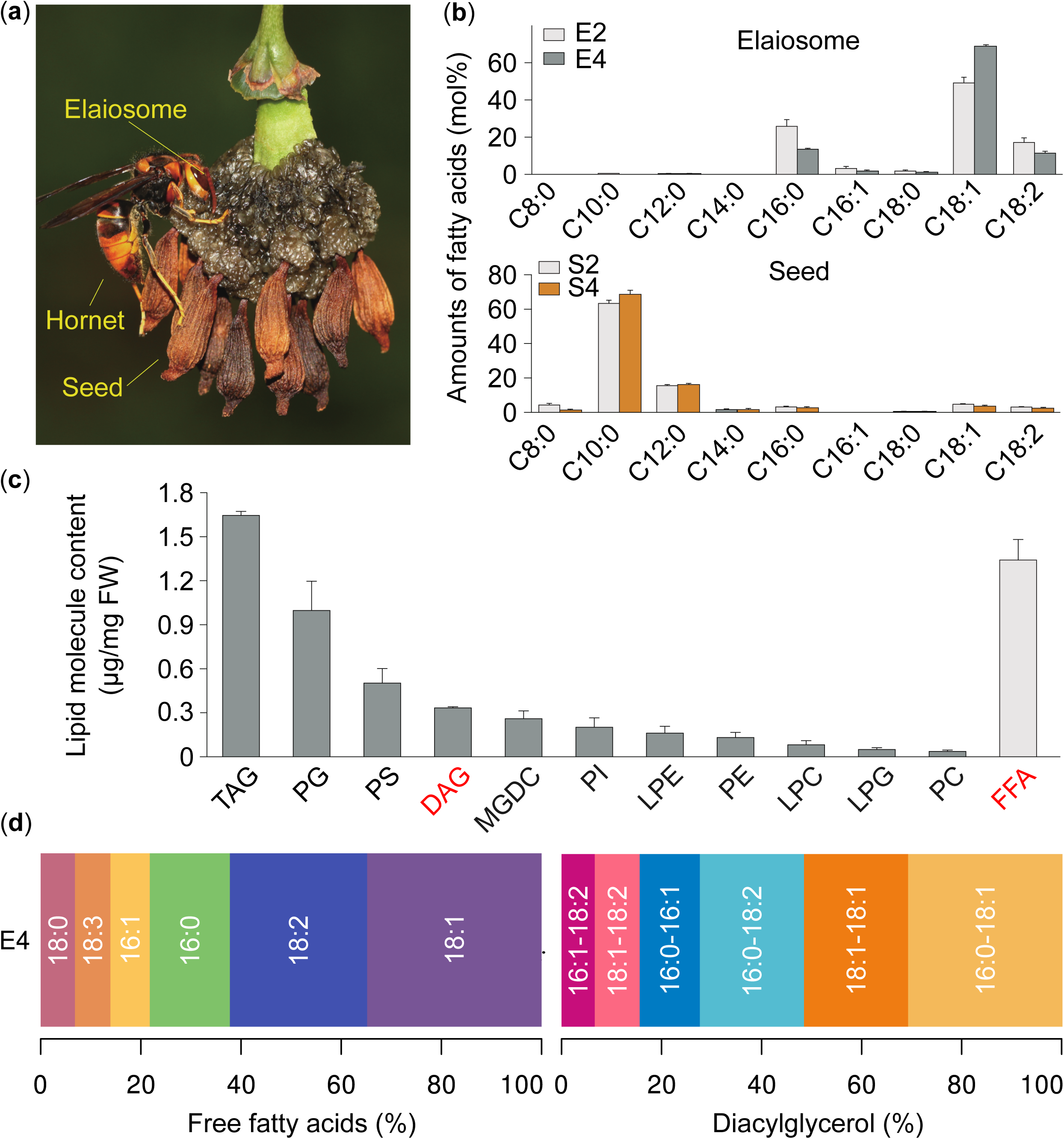
Wasp-mediated seed dispersal in *S. tuberosa* and lipids composition in elaiosome and seed. **(A)** *S. tuberosa* elaiosome being attacked by an individual of *Vespa velutina*. (**B**) Fatty acid composition extracted from all lipid molecules including free fatty acids in two developing elaiosomes (E2 and E4) and corresponding seeds (S2 and S4). **(C)** Lipid molecules including free fatty acids in mature elaiosome (E4) based on lipidomic analysis. Error bars showed the standard error across the three biological replicates. **(D)** the content percentage of different FFAs and DAG molecules in mature elaiosome (E4). Lipid molecules abbreviations are as follows: DAG, diacylglycerols; FFA, free fatty acids; LPC, lyso-phosphatidylcholines; LPE, lyso-phosphatidylethanolamines; LPG, Lyso-phosphatidylglycerol; MGDG, monogalactosyldiacylglycerol; PC, phosphatidylcholines; PE, phosphatidylethanolamines; PG, Phosphatidylglycerol; PI, phosphatidylinositol; PS, phosphatidylserine; TAG, triacylglycerols.

To quantify all lipid molecules (e.g. DAG) in mature elaiosomes (E4), we performed UPLC-QTOF/MS-based lipidomic analysis (*Materials and Methods*). Triacylglycerols (TAG) were the most abundant lipid class, followed by phosphatidylglycerol (PG) and phosphatidylserine (PS), with only small amounts of diacylglycerols (DAG) (Fig. 2c). As expected, oleic acid was the most abundant FFA, ∼35%, followed by C18:2 (∼27%) and C16:0 (∼16%) (Fig. 2d). Among DAGs, DAG (C16:0-C18:1) and 1,2-diolein (DAG[C18:1-C18:1]) were the major components, accounting for 31% and 21% of total DAGs, respectively (Fig. 2d). Thus, elaiosomes and seeds differ greatly in fatty acid composition, with lipid cues such as oleic acid and 1,2-diolein accumulating preferentially in elaiosomes.

### Expression patterns of lipid-related genes in seeds and elaiosomes of *S. tuberosa*

To dissect the molecular basis of the distinct fatty acid profiles of elaiosomes and seeds, we identified genes involved in fatty acid synthesis, glycerolipid assembly and oil body formation in *S. tuberosa* genome (Fig. 3a and Table S4). Transcriptome profiling across four stages of developing seeds (S1-S4) and corresponding elaiosome development (E1-E4) (Fig. S3) showed that 45 lipid-related genes were expressed in at least one tissue (FPKM ≥ 2) (Table S4). Most lipid-related genes exhibited high expression level in elaiosomes relative to seeds (Fig. 3b). Notably, genes involved in long-chain fatty acid biosynthesis, especially one *SAD* paralogue, were highly expressed in developing elaiosomes, while genes involved in medium-chain fatty acid synthesis (one FatB and two KASI) were specifically expressed in seeds, consistent with the contrasting fatty acid profiles of elaiosomes and seeds (Fig. 3b). We validated the expression pattern of key genes including *SAD*, *FatB* and *KASI* via RT-qPCR (Fig. 3c).

**Fig. 3.**
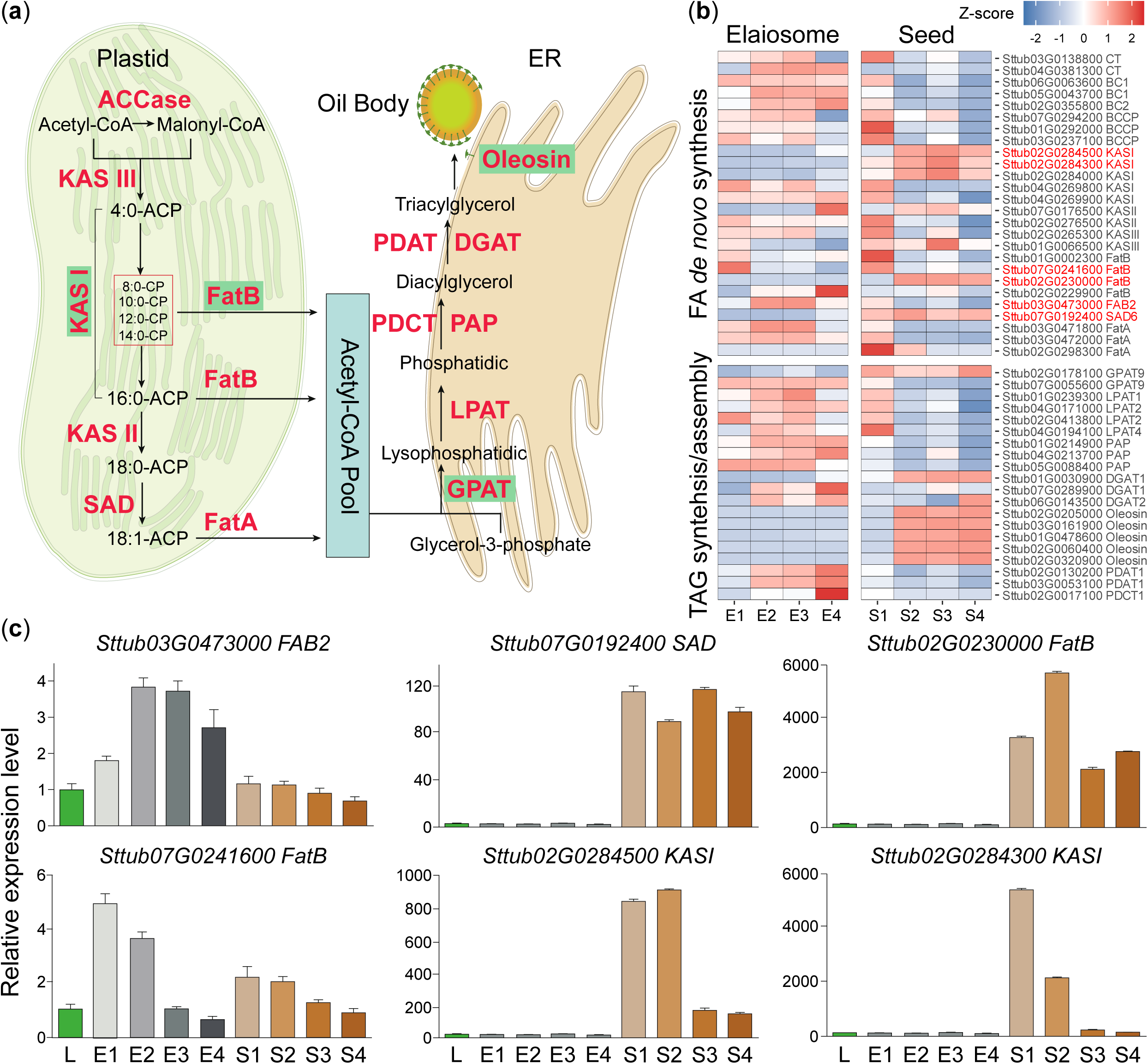
Genes involved into the biosynthesis of fatty acids and triacylglycerol (TAG) in *S. tuberosa*. **(A)** The biosynthetic pathway of fatty acids in plastid and triacylglycerols assembly in endoplasmic reticulum. The genes shaded in green show the expanded gene families in *S. tuberosa* genome. **(B)** Heatmap showing the expression profiles of lipid-related genes in *S. tuberosa* elaiosome and seed. The asterisks show the key lipid-genes for expression analysis. **(C)** Quantitative reverse transcription–polymerase chain reaction validation of gene expression we selected. Error bars showed the standard error across the three biological replicates, and the expression levels of gene in leaf sample (L) were normalized to 1.

### Functional analysis of *S. tuberosa* stearoyl-ACP Δ^9^ desaturases

Given the importance of oleic acid (C18:1) and its derivatives (e.g. 1,2-diolein) in insect-mediated seed dispersal (Skidmore & Heithaus, 1988; Brew *et al*., 1989; Lanza *et al*., 1992; Hughes *et al*., 1994; Pfeiffer *et al*., 2010; Boieiroa *et al*., 2012), we identified eight genes encoding stearoyl-ACPΔ^9^ desaturases (SADs) in the *S. tuberosa* genome (Table S4). Only two (*Sttub03G0473000.1* and *Sttub07G0192400.1*) were expressed in all tested tissues (Fig. 3c and Table S4). They shared high sequence identity with *Arabidopsis* SSI2/FAB2 (At2g43710, 68% amino acid sequence identity) and SAD6 (At1g43800, 66% identity), respectively, and were named *StFAB2* and *StSAD6*. *StFAB2* was highly expressed in developing elaiosomes (E1-E4), 5- to 7-fold higher than in developing seeds (S1-S4), whereas *StSAD6* was specifically expressed in developing seeds (Fig. 3b-c).

To characterize the potential function of *StFAB2* and *StSAD6*, we expressed each in the yeast BY4741 Δ*ole1* mutant, which is unable to synthesize unsaturated fatty acids and cannot survive without exogenous C18:1 (Fig. S4-S5, see *Materials and Methods*). The Δ*ole1* mutant showed a marked decrease in C16:1 (from 42.71% to 0.62%) and C18:1 (from 22.5% to 6.2%) relative to wild type (Fig. 4a-b). We heterologously overexpressed *StFAB2* and *StSAD6* in this background using an empty vector as negative control. Overexpression of *StFAB2* led to a 5-fold and 2.5-fold increase in C16:1 and C18:1, respectively (Fig. 4a-b), while *StSAD6* enhanced C16:1 and C18:1 by 1.9-fold and 1.6-fold (Fig. 4a-b). The content of C16:0 and C18:0 was slightly but not significantly reduced in both transformants. These results show that both enzymes can produce C16:1 and C18:1, with the greater accumulation in *StFAB2* transformants reflecting differences in intrinsic catalytic activity, protein abundance, or both. In elaiosomes, high *StFAB2* expression likely drives the accumulation of oleic acid, which can then be channelled into 1,2-diolein via the GPAT, LPAT, PAP and PDCT enzymes that are also highly expressed in this tissue (Fig. 3b).

**Fig. 4.**
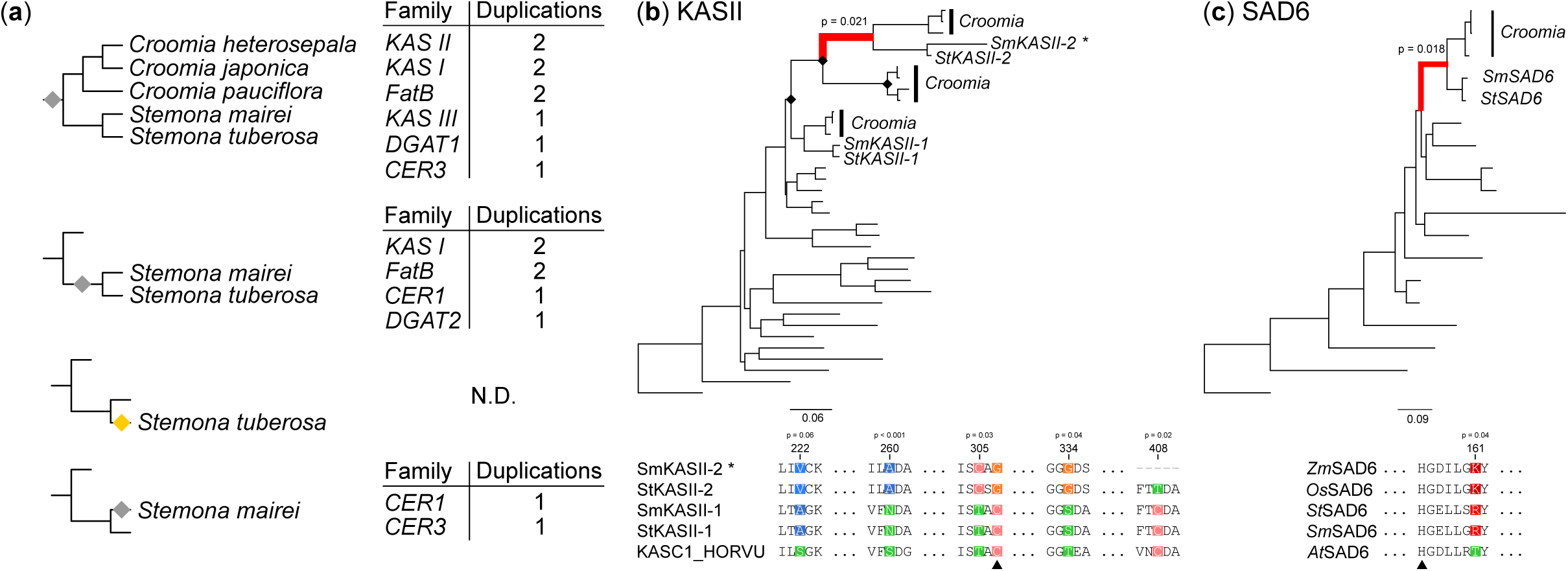
Functional characterization of *S. tuberosa* genes encoding SAD in yeast. (**A**) Long-chain fatty acid composition in wild-type yeast BY4741, *OLE1* mutant (*Δole1*), *Δole1* mutant with supplemented oleic acid, *Δole1* with over-expressed *StFAB2* or *StSAD6*. The fatty acid composition was identified by Gas Chromatography-Mass Spectrometry (GC-MS). Error bars showed the standard error across the five biological replicates. (**B**) GC-MS peaks showing the identified fatty acids in corresponding yeast described in Figure 4A. (**C**) Medium-chain fatty acids (MCFAs) analysed in the corresponding yeast mentioned in Figure 3A.

### Molecular basis for the accumulation of MCFAs in *S. tuberosa* seed

Oleic acid and derivatives (e.g. 1,2-diolein and hydrocarbons) are the key functional compounds in elaiosomes, but MCFAs are frequently found in seeds dispersed by insects (Lanza *et al*., 1992; Hughes *et al*., 1994; Fischer *et al*., 2008; Chen *et al*., 2016), suggesting a resource trade-off between elaiosomes and seeds. However, the molecular mechanisms underlying this resource allocation remain elusive.

Intriguingly, we observed a marked increase in medium-chain fatty acids in the Δ*ole1* mutant, including C10:0, C12:0 and C14:0 (17.5-fold, 3.7-fold and 3.8-fold relative to wild type) (Fig. 4b-c). Conversely, when Δole1 was grown on agar plates with exogenous oleic acid, MCFA levels were negligible (Fig. 4b-c). This indicated a compensatory mechanism between oleic acid and MCFAs in yeast, likely reflecting the use of shorter-chain fatty acids to maintain membrane fluidity in the absence of unsaturated fatty acids. In *S. tuberosa* seeds, the dedicated *FatB* thioesterase machinery described below may reinforce this metabolic relationship, actively channelling flux toward MCFAs.

One FatB (*Sttub02G0230000.1*) was specifically and highly expressed in seeds (Fig. 3). Plant FatB thioesterases are known to play a critical role in MCFA production (Kalinger et al., 2020). We overexpressed all four *StFatB* genes in the yeast strain INVSc1, which does not produce capric acid (C10:0) (Fig. 5a). Only *StFatB* (*Sttub02G0230000.1*) produced C10:0 in yeast and significantly enhanced C12:0 and C14:0 levels (Fig. 5a). Heterologous expression of the same *StFatB* in Arabidopsis also led to significant enrichment of MCFAs in transgenic seeds (Fig. 5b). These results indicate that *StFatB* is sufficient to drive MCFA synthesis.

**Fig. 5.**
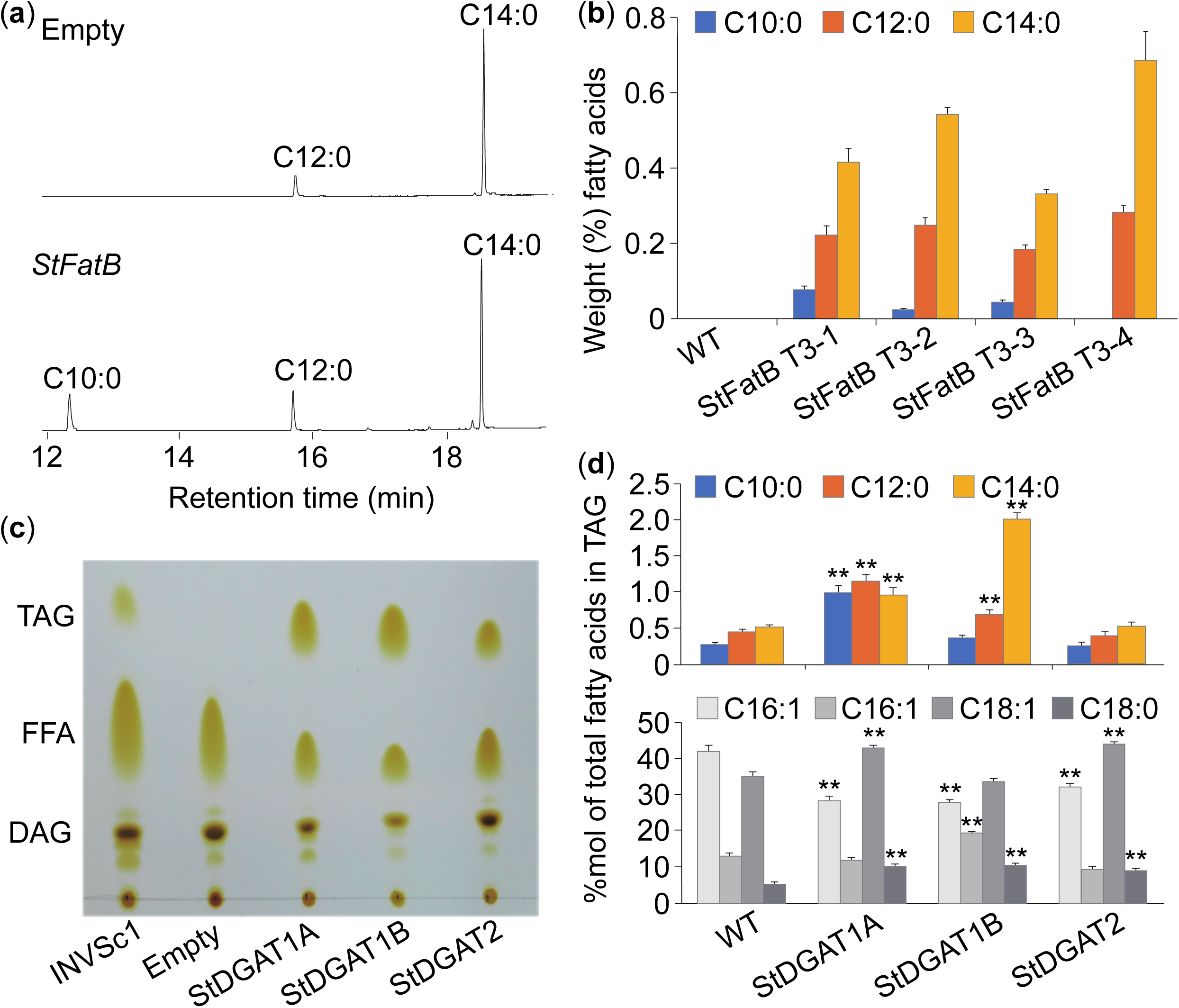
Functional characterization of *S. tuberosa* genes encoding FatB and DGATs. (**A**) GC-MS peaks showing the medium-chain fatty acids in wild-type yeast INVSc1 and yeast with over-expressed *StFatB.* **(B)** Relative proportions of C10:0, C12:0 and C14:0 in seeds of *A. thaliana* wild-type (Col-0) and four T3 homozygous transgenic lines expressing the *StFatB* gene under control of the 35S promoter. Error bars indicate ±SE (n = 3). **(C)** Overexpression of *StDGAT1A/B* and *StDGAT2* restores TAG biosynthesis in TAG-deficient yeast mutant strain H1246 as visualized in TLC. Triacylglycerol (TAG), diacylglycerol (DAG) and free fatty acid (FFA) were separated in TLC plate and visualized by iodine staining. H1246 transformed with the empty pYES2 vector was used as negative control and wild type yeast INVSc1 served as a positive control. **(D)** FA composition of TAG product isolated from H1246 yeast cells expressing *StDGAT1A/B* and *StDGAT2*. Wild type yeast INVSc1 was used as control. Error bars indicate ±SE (n = 3). Asterisks (**) indicate the *p*-value less than 0.01 (student t-test).

However, MCFA levels achieved in transgenic yeast and Arabidopsis seeds were substantially lower than those in *S. tuberosa* seeds, suggesting that efficient incorporation of MCFAs into triacylglycerol (TAG) requires additional enzymes. Diacylglycerol acyltransferases (DGAT) are known to play a critical role in the accumulation of unusual fatty acids in TAG (Turchetto-Zolet et al., 2016). We identified three DGAT genes: two *DGAT1* paralogues, *StDGAT1A* (Sttub01G0030900.1) and *StDGAT1B* (Sttub07G0289900.1), and *StDGAT2* (Sttub06G0143500.1). *StDGAT1A* showed high expression in developing seeds relative to elaiosomes, while *StDGAT1B* showed low expression in all tested tissues (Fig. 3b). *StDGAT2* also showed higher expression in developing seeds than elaiosomes (Fig. 3b). We confirmed TAG biosynthesis by heterologous expression in the TAG-deficient yeast strain H1246 (see *Materials and Methods*). TLC analysis showed that transformants overexpressing *StDGAT1A/B* and *StDGAT2* restored TAG biosynthesis (Fig. 5c). To determine the substrate preference of these DGATs, we performed yeast feeding assays. Feeding exogenous C8:0, C10:0 and C12:0 to transformants expressing *StDGAT1A/B* and *StDGAT2* revealed distinctive substrate preferences (Fig. 5d). *StDGAT1A* showed substrate specificity toward MCFAs and C18:1, while *StDGAT1B* preferred C12:0, C14:0 and C16:0 (Fig. 5d). *StDGAT2* showed no activity on MCFAs but was active on C18:1 (Fig. 5d). None of the three DGATs showed activity on C8:0 (Fig. 5d). Overall, seed-specific *StFatB* has a major contribution to MCFA production in *S. tuberosa* seeds, with MCFAs preferentially incorporated into TAG via *StDGAT1* paralogues.

### Hydrocarbons biosynthesis in *S. tuberosa* elaiosomes

Fatty acid-derived hydrocarbons (alkanes and alkenes) in elaiosomes of *S. tuberosa* differ significantly from those in leaves, and are critical for wasp-mediated seed dispersal (Chen *et al*., 2018a). (Z)-9-tricosene (C23:1Δ9), a pheromone component produced by various insects (Carlson *et al*., 1971), is considered the primary attractant for wasps (Chen *et al*., 2018a), suggesting that *S. tuberosa* employs chemical mimicry to elicit dispersal behaviour. Plant hydrocarbons are typically synthesized through elongation of fatty acyl-CoA (e.g. C16:0, C16:1, C18:0 and C18:1) into very-long-chain fatty acyl-CoA, which are then reduced to aldehydes and decarbonylated to yield alkanes or alkenes with one fewer carbon (Kang & Nielsen, 2017; Fig. 6a). (Z)-9-tricosene is accordingly derived from C18:1Δ9-CoA (Fig. 6a), and the large amounts of free C18:1 in elaiosomes (Fig. 2d) provide an abundant precursor. CER1 and CER3 (ECERIFERUM), encoding fatty acyl decarbonylase and reductase, respectively, are core components of hydrocarbon biosynthesis in Arabidopsis (Aarts *et al*., 1995; Bourdenx *et al*., 2011; Bernard *et al*., 2012). We identified single-copy genes encoding *CER1* (*Sttub01G0253000.1*) and *CER3* (*Sttub06G0047800.1*) in *S. tuberosa*. Both exhibited highly coordinated, elaiosome-specific expression (Pearson’s R = 0.99; Fig. 6b), although their role in hydrocarbon biosynthesis remains to be functionally validated.

**Fig. 6.**
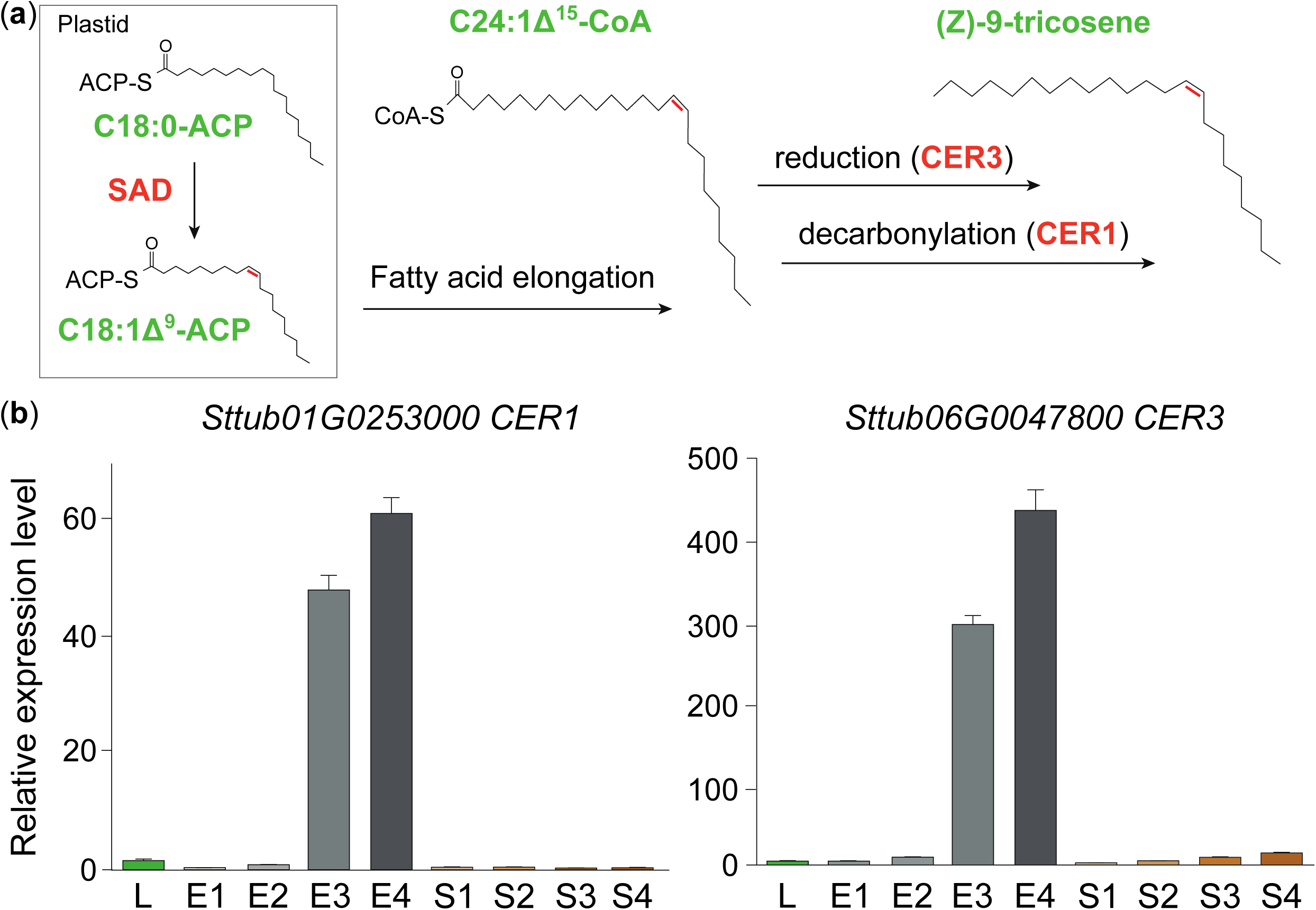
Putative biosynthetic pathway for (*Z*)-9-tricosene, the key compound that recruits wasp dispersers, and key genes in *S. tuberosa*. (**A**) Proposed biosynthetic pathway and major reaction of (*Z*)-9-tricosene (C23:1Δ^9^ alkene). (**B**) Validation of expression pattern of *S. tuberosa CER1* and *CER3* via quantitative reverse transcription–polymerase chain reaction. Error bars show the standard error across the three biological replicates, and the expression levels of gene in the leaf sample (L) were normalized to 1.

### Diversifying selection on fatty acid synthesis genes during the evolution of myrmecochory

To test whether the transition from myrmecochory to vespicochory involved changes in lipid-related gene copy number, we inferred phylogenies for the main lipid-related genes expressed in *S. tuberosa* elaiosomes and seeds. Several gene families showed lineage-specific expansion in *Stemona*, including KAS I (carbon chain elongation, C4 to C16), FatB (release of medium-chain acyl groups, C8-C14), GPAT (rate-limiting step in de novo glycerolipid synthesis) and oleosin (lipid droplet packaging) (Figs. 3, S6). Gene trees indicated that most expansions occurred along the branch subtending *Stemona*, yielding paralogues shared by *S. tuberosa* and *S. mairei*, while a smaller number of species-specific duplications and losses produced copy number differences, notably in KAS I, PDCT1, CER1 and CER3 (Fig. S6).

We then employed codon models to search for evidence of diversifying selection on fatty acid synthesis genes during the transition to myrmecochory and vespicochory. We fitted ABSREL models to test for episodic diversifying selection along three sets of branches: those ancestral to *Croomia* + *Stemona*; those subtending *Stemona*; and those leading to *S. tuberosa*. Filtering out positive results originating from alignment errors, we recovered robust signals for episodic diversifying selection along the branch ancestral to *Croomia* and *Stemona* in KASII and SAD6 (Fig. 7). In KASII, this branch occurs following an inferred gene duplication in *Stemonaceae* (Fig. 7a; Fig. S6); notably, the resulting *S. mairei* copy is partially truncated, a result not obviously due to annotation error. MEME analysis revealed evidence for diversifying selection at five individual sites (Fig. 7b). One was adjacent to a position with an inferred cysteine to glycine substitution, annotated in the active site for beta-ketoacyl synthase activity based on KASC1_HORVU (Fig. 7b). In *SAD6*, the signal occurs along a branch subtending *Stemonaceae* orthologues, although it is the daughter of a branch that conflicts with the species tree, albeit with low support (Fig. 7c; Fig. S6). MEME identified only a single site, containing an inferred lysine to arginine substitution, homologous to a position seven amino acids distant from a Fe cation-coordinating histidine in *At*SAD6 (Fig. 7d).

**Figure 7.**
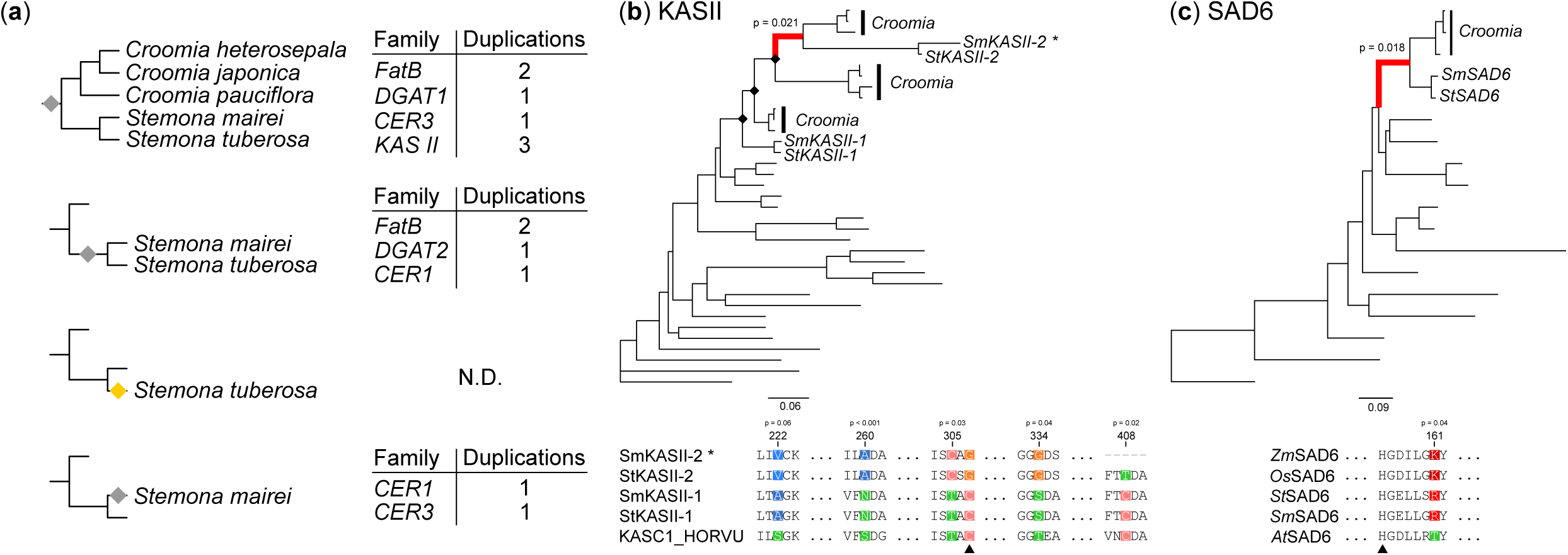
Diversifying selection in fatty acid synthesis genes. Results from ABSREL tests for episodic diversifying selection. Red branches show tested foreground branches that were significant at the 5% level. Diamonds at nodes indicate inferred gene duplications based on species overlap. Aligned sequences show results for MEME tests for site-level diversifying selection along the inferred branch. Positions are relative to *St*KASII-1 and *St*SAD6. P-values show the results from the MEME test for diversifying selection. (a, b) KASII. Black arrow shows the position of an active site residue in *Hordeum vulgare* KASCI. (d, e) SAD6. Black arrow shows the position of a Fe cofactor coordinating histidine in the active site of *Arabidopsis thaliana* SAD6.

## Discussion

### Molecular basis of elaiosome lipid cues in *Stemona tuberosa*

Over 11,000 plant species have evolved elaiosomes as a reward to ants that disperse seeds (Lengyel *et al*., 2010). Lipid composition analyses of elaiosomes from different plant groups have highlighted the importance of oleic acid and 1,2-diolein in insect-mediated seed dispersal, suggesting strong convergent evolution (Lanza *et al*., 1992; Hughes *et al*., 1994; Fischer *et al*., 2008). The lipid composition of elaiosomes was thought to resemble that of insect prey rather than seeds, particularly 1,2-diolein, which is an important component of insect haemolymph (Hughes *et al*., 1994). However, the molecular basis underlying these lipid cues have remained unclear.

*Stemona tuberosa* develops unusually large elaiosomes that serve as rewards offered to wasps (vespicochory) to disperse seeds (Chen *et al*., 2018a). Combining genomic, transcriptomic, lipidomic and functional analyses, we showed that high expression of one SAD paralogue (stearoyl-ACP Δ9 desaturase) in *S. tuberosa* elaiosomes drives the production of oleic acid (C18:1), which is channelled into 1,2-diolein through the glycerolipid assembly pathway, consistent with the high expression of GPAT, LPAT and PAP in elaiosomes (Fig. 3b). Genes encoding DGAT and oleosin were expressed at low levels during elaiosome development but were predominantly expressed in developing seeds (Fig. 3b and Table S4), implying that oleic acid and 1,2-diolein are not efficiently incorporated into triacylglycerol (TAG) or packaged into lipid droplets. This is consistent with the large amounts of free oleic acid and 1,2-diolein detected in *S. tuberosa* elaiosomes (Fig. 2d). These free lipid molecules are likely critical for attracting wasps and promoting seed dispersal. In addition, the abundant TAGs and free polyunsaturated fatty acids (e.g. C18:2, C18:3) detected in elaiosomes may serve as energy and nutrients rewarding seed dispersers (Hughes *et al*., 1994). Given the convergent evolution of lipid cues across elaiosome-bearing plants (Fischer *et al*., 2008; Pfeiffer *et al*., 2010; Boieiroa *et al*., 2012), our results provide a framework for investigating whether similar biosynthetic pathways underlie elaiosome lipid cues in other lineages (Fig. 8).

**Fig. 8.**
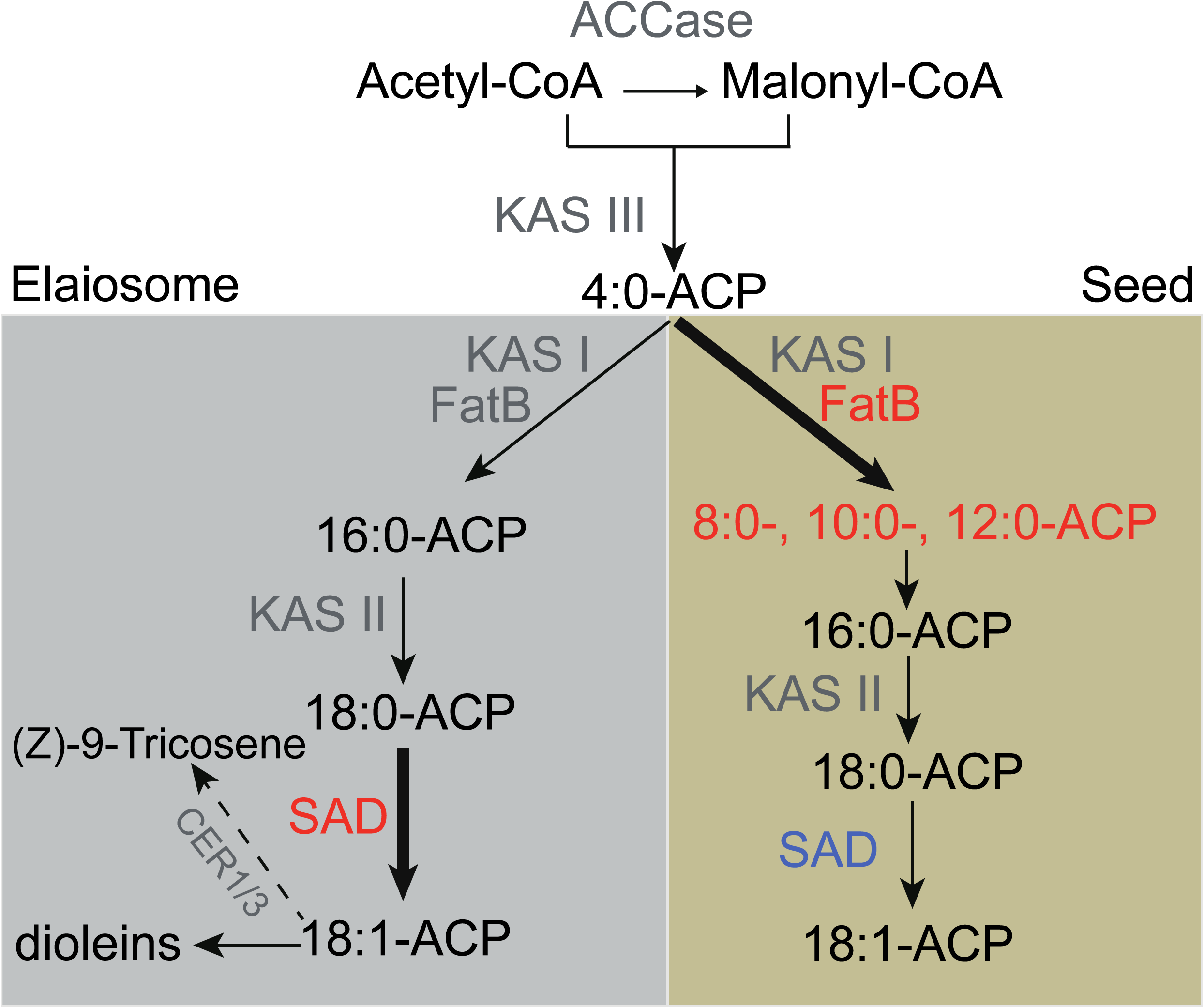
Proposed model explaining the differentiate fatty acid species in *S. tuberosa* elaiosome (grey) and seed (brown). The high expression of one of *SAD* genes results in the accumulation of oleic acids (C18:1) and its derivatives (e.g. 1,2-dioleins and (Z)-9-tricosene) in elaiosomes, promoting seed dispersal. On the contrary, gene expansion and seed-specific expression of *FatB* result in large amounts of MCFAs and low level of oleic acids in seeds. Red and blue fonts represent genes that are expressed at a high versus low level, respectively. Thick black arrows indicate the major reaction of fatty acids biosynthesis.

### Hydrocarbon biosynthesis and chemical mimicry

Beyond these conserved lipid cues, fatty acid-derived hydrocarbons specifically synthesized in the mature *S. tuberosa* elaiosome attract hornets and promote seed dispersal efficiency (Chen *et al*., 2018a). Among these, (Z)-9-tricosene, the principal wasp attractant in *S. tuberosa* (Chen *et al*., 2018a), is also a major pheromone component of various female insects (Carlson et al., 1971). A parallel is found in sexually deceptive orchids, where organ-specific expression of SAD genes mediates the production of pollinator sex pheromones (Schlüter *et al*., 2011; Sedeek *et al*., 2016). In *S. tuberosa*, the high elaiosome-specific expression of one SAD gene produces abundant free oleic acid, providing the precursor for (Z)-9-tricosene biosynthesis. We also identified two elaiosome-specific genes, *CER1* and *CER3*, which are core components of the hydrocarbon biosynthesis pathway (Aarts *et al*., 1995; Bourdenx *et al*., 2011; Bernard *et al*., 2012) and may be key in the biosynthesis of (Z)-9-tricosene. Notably, in *Aquilaria sinensis* (Thymelaeaceae), the only lineage where vespicochory evolved *de novo* rather than from myrmecochorous ancestors, short-carbon-chain (C5-C9) volatiles serve as the main attractants, and their rapid diffusion is thought to facilitate dispersal of short-lived seeds (Qin *et al*., 2022). Together, these findings indicate that fatty acid metabolism provides a versatile biochemical platform for insect-mediated seed dispersal, with SAD-derived oleic acid serving both as a conserved lipid cue and as a precursor for lineage-specific recruitment compounds.

### Resource trade-off and medium-chain fatty acid accumulation in seeds

The evolution of elaiosomes to enhance seed dispersal likely entails a resource trade-off between seeds and elaiosomes, since resources invested in lipid-rich elaiosomes could alternatively be allocated to seeds. Seeds and elaiosomes from different plant species differ significantly in chemical composition, particularly fatty acid profiles (Lanza *et al*., 1992; Hughes *et al*., 1994; Fischer *et al*., 2008), and this difference is especially pronounced in *S. tuberosa*, which develops large elaiosomes and shows striking differences in fatty acid composition between seeds and elaiosomes (Fig. 2). We found that genes encoding acyl-ACP thioesterase B (*FatB*) have expanded significantly in the *S. tuberosa* genome. *FatB* enzymes are considered major determinants of fatty acid carbon chain length, with substrate preference for C8:0-C18:0 (Jones *et al*., 1995), and specialized *FatB* variants with substrate specificity for MCFAs have been identified in several plant species (Voelker *et al*., 1992; Dehesh *et al*., 1996; Jing *et al*., 2011). We identified a seed-specific FatB whose overexpression in yeast and Arabidopsis seeds increased MCFA accumulation (Fig. 5a-b). Notably, although SAD genes were expressed in developing *S. tuberosa* seeds and had the capacity to produce oleic acid (Fig. 4a-b), only trace amounts of oleic acid were detected in seeds (Fig. 2b). We speculate that the high, seed-specific expression of *FatB* releases acyl chains at medium chain lengths, reducing the pool of C18:0-ACP substrate available for desaturation to oleic acid (Fig. 8). Furthermore, StDGAT1A and StDGAT1B exhibited substrate preference for MCFAs, suggesting that DGAT1 substrate specificity evolved in concert with *FatB*-mediated MCFA production in *S. tuberosa* seeds, paralleling the coordinated evolution of acyltransferase specificity and unusual fatty acid accumulation reported in other plants (Burgal *et al*., 2008; Horn *et al*., 2016).

### Comparative genomics points to a shared genetic basis for myrmecochory and vespicochory

To determine whether the functional changes we identified are specific to vespicochory or reflect a more general role in insect-mediated seed dispersal, we turned to comparative analysis with *Stemona mairei*, an ant-dispersed species that also produces elaiosomes. Many of the lineage-specific gene family expansions in fatty acid synthesis genes are shared between the two species, and significant signals of episodic diversifying selection were limited to branches subtending *Stemona* and *Croomia*, a related genus of ant-dispersed, elaiosome-bearing species (Fig. 1b). Although comparative evidence for specific sequence-level functional changes is limited, this pattern suggests that such changes may have occurred during the transition to myrmecochory rather than during the later specialization to vespicochory in *S. tuberosa*. One of two elaiosome-specific *CER* genes detected in *S. tuberosa* have duplicated independently in *S. mairei*, suggesting that their role in mediating wasp attraction may be more complex than a simple gain of function, or governed primarily by expression differences. Further comparative expression analysis and functional characterization across the two species will be necessary to resolve this. *Stemona* represents a promising system because of repeated transitions between myrmecochory and vespicochory, which could provide additional power to detect convergent signals in broader comparisons across the genus.

Our two new *Stemona* genomes reveal the molecular architecture underlying fatty acid differentiation between elaiosomes and seeds, and provide the first genomic framework linking lipid biosynthesis to insect-mediated seed dispersal. Because oleic acid and 1,2-diolein are key cues in both myrmecochory and vespicochory, the shared biochemical basis documented here may help explain how vespicochory repeatedly evolves from ant-dispersed ancestors. More broadly, these findings pave the way to comparative studies across elaiosome-bearing lineages and to investigations of the small genetic and regulatory changes that tip the balance from ant to wasp dispersal.

## Author contributions

Conceptualization: GChen, GC; Investigation: WX, TY, NWH, CZ, ZH, GChen; Formal Analysis: WX, TY, NWH, CZ, ZH; Visualization: NWH, WX, TY, CZ, ZH; Funding acquisition: GChen; Writing original draft: G.Chen, GC; Writing – review & editing: all co-authors.

## Abbreviations

ACCase: Acetyl-CoA carboxylase
bp: base pair
BUSCO: Benchmarking Universal Single-Copy Orthologs
DEG: differentially expressed gene
DGAT: Diglyceride acyltransferase
FatB: acyl-ACP thioesterases B
Gb: giga-base
GO: Gene Ontology
GPAT: glycerol-3-phosphate acyltransferase
KASI: ketoacyl-acyl carrier protein synthase I
KASIII: ketoacyl-acyl carrier protein synthase III
kb: kilobase
KEGG: Kyoto Encyclopedia of Genes and Genomes
LPAT: Lysophosphatidylglycerol acyltransferase
LTR: long terminal repeat
Mbp: megabase pair
MCFAs: medium-chain fatty acids
MRCA: most recent common ancestor
NCBI: The National Center for Biotechnology Information
PAP: Polyphosphate:AMP phosphotransferase
PDAT: phospholipid:diacylglycerol acyltransferase
PDCT: phosphatidylcholine diacylglycerol cholinephosphotransferase
qRT-PCR: quantitative real-time reverse-transcription polymerase chain reaction
SAD: stearoyl-ACP Δ^9^ desaturases.

## Data availability

The genome sequencing data and transcriptome data generated in this study are available via NCBI under the BioProject accessions PRJNA985082 and PRJNA985114, respectively.

## Supporting Information

Additional Supporting Information may be found online in the Supporting Information section at the end of the article.

**Figure S1.** GenomeScope 2.0 k-mer frequency profiles for (A) *Stemona tuberosa* and (B) *S. mairei*, based on Illumina short-read data with k = 19 and ploidy = 2. The x-axis shows k-mer coverage depth and the y-axis shows coverage x frequency. The observed k-mer spectrum (blue), fitted model (black), unique sequence component (yellow), and error component (orange) are shown; dashed vertical lines mark the heterozygous and homozygous k-mer coverage peaks. Estimated genome sizes were 899.8 Mb for S. tuberosa (heterozygosity 0.401%, error rate 0.013%, duplication rate 1.18) and 951.7 Mb for *S. mairei* (heterozygosity 1.73%, error rate 0.126%, duplication rate 4.26). The more prominent heterozygous peak and higher duplication rate in *S. mairei* are consistent with its larger genome and greater repeat content.

**Figure S2.** Hi-C map (High-throughput chromosome conformation capture) of the *Stemona tuberosa* (a) and *S. mairei* (b) genomes showing strong interactive signal within intra-chromosome and at diagonal regions. The color bar indicates the interactive signal.

**Figure S3.** The shape, size and colors of seed (S1-S4) and elaiosome (E1-E4) during fruit development and ripening at different stages. Bar, 1 cm.

**Figure S4.** Live/dead spot assay of yeast BY4741 mutant strain *Δole1* (*MATa ole1Δ::HIS3 leu2Δ0 met15Δ0 ura3Δ0*) on synthetic complete selection medium (-His) with (right panel) or without (left panel) oleic acid supplement.

**Figure S5.** Construction and molecular identification of *OLE1* mutant (*Δole1*) in yeast strain BY4741. (A). Schematic representation of *OLE1* genomic DNA from ATG to TAA. The black line in the blue box shows the region that will be disrupted. (B). Schematic representation of *OLE1* deletion cassette. The left and right arm (light blue box) are homologous regions of *OLE1* genomic DNA as shown in A. The light green box represents the HIS3 expression cassette. Specific primers used for construction of *OLE1* deletion cassette and molecular identification for yeast strain BY474141 and *Δole1* are indicated by the arrows. (C). Polymerase chain reaction (PCR)-based identification of *Δole1* mutant line with specific primers.

**Figure S6.** Gene trees of fatty acid synthesis gene families. a-l) gene trees of each major family discussed in text. Diamonds at nodes indicate inferred gene duplications based on species overlaps. Scale bars give the expected number of nucleotide substitutions per site. Branch labels are support values from the SH-aLRT/Ultrafast Bootstrap.

## Supplementary tables

**Table S1.** Public RNA seq datasets used for annotation in this study.

**Table S2.** Genomes and transcriptomes used in this study.

**Table S3.** Primer information used in this study.

**Table S4.** Expression of genes involved into fatty acids synthesis, glycerolipid assembly and oil body formation in *Stemona tuberosa*. E1-E4 are elaiosome stages and S1-S4 are seed stages. Values are FPKM.

**Table S5.** Repeat annotation.

**Table S6**. Sequencing coverage, assembly and annotation summary.

**Table S7**. BUSCO assessment.

**Table S8**. Functional annotation.

**Table S9**. Gene family expansions and contractions inferred in *S. tuberosa*.

**Table S10**. Gene family expansions and contractions inferred in the *S. tuberosa-S. mairei* branch.

## References

Aarts MG, Keijzer CJ, Stiekema WJ, Pereira A. 1995. Molecular characterization of the *CER1* gene of *Arabidopsis* involved in epicuticular wax biosynthesis and pollen fertility. Plant Cell 7: 2115–2127.

Karnish A. 2024. Seed Dispersal by Ants: A Primer. International Journal of Plant Sciences 185: 403–411.

Bernard A, Domergue F, Pascal S, Jetter R, Renne C, Faure JD, Haslam RP, Napier JA, Lessire R, Joubès J. 2012. Reconstitution of plant alkane biosynthesis in yeast demonstrates that Arabidopsis ECERIFERUM1 and ECERIFERUM3 are core components of a very-long-chain alkane synthesis complex. Plant Cell 24: 3106–3118.

Boieiroa M, Espadaler X, Gómez C, Eustaquio, A. 2012. Spatial variation in the fatty acid composition of elaiosomes in an ant-dispersed plant: differences within and between individuals and populations. Flora 207: 497–502.

Bourdenx B, et al. 2011. Overexpression of *Arabidopsis ECERIFERUM1* promotes wax very-long-chain alkane biosynthesis and influences plant response to biotic and abiotic stresses. Plant Physiology 156: 29–45.

Brew CR, O’Dowd DJ, Rae ID. 1989. Seed dispersal by ants: behaviour-releasing compounds in elaiosomes. Oecologia 80: 490–497.

Brown JW, Walker JF, Smith SA. 2017. Phyx: phylogenetic tools for unix. Bioinformatics 33: 1886–1888.

Buchfink B, Reuter K, Drost H-G. 2021. Sensitive protein alignments at tree-of-life scale using DIAMOND. Nature Methods 18: 366–368.

Burgal J, Shockey J, Lu C, Dyer J, Larson T, Graham I, Browse J. 2008. Metabolic engineering of hydroxy fatty acid production in plants: RcDGAT2 drives dramatic increases in ricinoleate levels in seed oil. Plant Biotechnology Journal 6: 819–831.

Burton JN, et al. 2013. Chromosome-scale scaffolding of de novo genome assemblies based on chromatin interactions. Nature Biotechnology 31: 1119–1125.

Carlson DA, et al. 1971. Sex attractant pheromone of the house fly: isolation, identification and synthesis. Science. 174, 76–78 (1971).

Chen G, Huang SZ, Chen SC, Chen YH, Liu X, Sun WB. 2016. Chemical composition of diaspores of the myrmecochorous plant *Stemona tuberosa* Lour. *Biochem*. Biochemical Systematics and Ecology 64: 31–37.

Chen G, Wang ZW, Wen P, Wei W, Chen Y, Ai H, Sun WB. 2018a. Hydrocarbons mediate seed dispersal: a new mechanism of vespicochory. New Phytologist 220: 714–725.

Chen S, Zhou Y, Chen Y, Gu J. 2018b. fastp: an ultra-fast all-in-one FASTQ preprocessor. Bioinformatics 34: i884–i890.

Chen YS, Zeng CX, Muellner-Riehl AN, Wang ZH, Sun L, Schinnerl J, Kongkiatpaiboon S, Kadota Y, Cai XH, Chen G. 2021. Invertebrate-mediated dispersal plays an important role in shaping the current distribution of a herbaceous monocot. Journal of Biogeography 48: 1101–1111.

Ciccarelli D, Andreucci AC, Pagni AM, Garbari F. 2005. Structure and development of the elaiosome in *Myrtus communis* L. (Myrtaceae) seeds. Flora, 200, 326–331.

Clough SJ, Bent AF. 1998. Floral dip: a simplified method for Agrobacterium-mediated transformation of *Arabidopsis thaliana*. Plant Journal 16: 735–743.

Dehesh K, Jones A, Knutzon DS, Voelker TA. 1996. Production of high levels of 8:0 and 10:0 fatty acids in transgenic canola by overexpression of *ChFatB2*, a thioesterase cDNA from *Cuphea hookeriana*. Plant Journal 9: 167–172.

dos Reis M, Yang Z. 2011. Approximate Likelihood Calculation on a Phylogeny for Bayesian Estimation of Divergence Times. Molecular Biology and Evolution 28: 2161–2172.

Edwards W, Dunlop M, Rodgerson L. 2006. The evolution of rewards: seed dispersal, seed size and elaiosome size. Journal of Ecology 94: 687–694.

Farwig N, Berens DG. 2012. Imagine a world without seed dispersers: a review of threats, consequences and future directions. Basic and Applied Ecology 13: 109–115.

Fischer RC, Richter A, Hadacek F, Mayer V. 2008. Chemical differences between seeds and elaiosomes indicate an adaptation to nutritional needs of ants. Oecologia 155: 539–547.

Fu L, Niu B, Zhu Z, Wu S, Li W. 2012. CD-HIT: accelerated for clustering the next-generation sequencing data. Bioinformatics 28: 3150–3152.

Gabriel L, Brůna T, Hoff KJ, Ebel M, Lomsadze A, Borodovsky M, Stanke M. 2024. BRAKER3: Fully automated genome annotation using RNA-seq and protein evidence with GeneMark-ETP, AUGUSTUS, and TSEBRA. Genome Research 34: 769–777.

Gietz RD, Schiestl RH. 2007. High-efficiency yeast transformation using the LiAc/SS carrier DNA/PEG method. Nature Protocols 2: 31–34.

Grabherr MG, Haas BJ, Yassour M, Levin JZ, Thompson DA, Amit I, Adiconis X, Fan L, Raychowdhury R, Zeng Q, et al. 2011. Full-length transcriptome assembly from RNA-Seq data without a reference genome. Nature Biotechnology 29: 644–652.

Greger H. 2019. Structural classification and biological activities of *Stemona* alkaloids. Phytochemistry Review 18: 463–493.

Haas BJ, et al. 2008. Automated eukaryotic gene structure annotation using EVidenceModeler and the Program to Assemble Spliced Alignments. Genome Biology 9: R7.

Holt C, Yandell M. 2011. MAKER2: an annotation pipeline and genome-database management tool for second-generation genome projects. BMC Bioinformatics 12: 491.

Horn PJ, Liu J, Cocuron JC, McGlew K, Thrower NA, Larson M, Lu C, Alonso AP, Ohlrogge J. 2016. Identification of multiple lipid genes with modifications in expression and sequence associated with the evolution of hydroxy fatty acid accumulation in *Physaria fendleri*. Plant Journal 86: 322–348.

Howe HF, Smallwood J. 1982. Ecology of seed dispersal. Annual review of ecology, evolution and systematics 13: 201–228.

Huang L, Gao L, Chen C. 2021. Role of Medium-Chain Fatty Acids in Healthy Metabolism: A Clinical Perspective. Trends in Endocrinology and Metabolism 32: 351–366.

Hughes L, Westoby M, Jurado E. 1994. Convergence of elaiosomes and insect prey: evidence from ant foraging behaviour and fatty acid composition. Functional Ecology 8: 358–365.

Janzen DH. 1970. Herbivores and the number of tree species in tropical forests. American Naturalist 104: 501–528.

Jing F, Cantu DC, Tvaruzkova J, Chipman JP, Nikolau BJ, Yandeau-Nelson MD, Reilly PJ. 2011. Phylogenetic and experimental characterization of an acyl-ACP thioesterase family reveals significant diversity in enzymatic specificity and activity. BMC Biochemistry 12: 44.

Jones, A., Davies, H.M., Voelker, T., 1995. Palmitoyl-acyl carrier protein (ACP) thioesterase and the evolutionary origin of plant acyl-ACP thioesterases. Plant Cell 7: 359–371.

Jordano P, García C, Godoy JA, Garcia-Castaño J. 2007. Differential contribution of frugivores to complex seed dispersal patterns. Proceedings of the National Academy of Sciences of the U.S.A. 104: 3278–3282.

Kalinger RS, Pulsifer IP, Hepworth SR, Rowland O. 2020. Fatty Acyl Synthetases and Thioesterases in Plant Lipid Metabolism: Diverse Functions and Biotechnological Applications. Lipids 55:435–455.

Kang MK, Nielsen J. 2017. Biobased production of alkanes and alkenes through metabolic engineering of microorganisms. Journal of Industrial Microbiology and Biotechnology 44: 613–622.

Karnish, A. 2024. Seed Dispersal by Ants: A Primer. International Journal of Plant Sciences 185: 403–411.

Katoh K, Standley DM. 2013. MAFFT Multiple Sequence Alignment Software Version 7: Improvements in Performance and Usability. Molecular Biology and Evolution 30: 772–780.

Kim D, Paggi JM, Park C, Bennett C, Salzberg SL. 2019. Graph-based genome alignment and genotyping with HISAT2 and HISAT-genotype. Nature Biotechnology 37: 907–915.

Korf I. 2004. Gene finding in novel genomes. BMC Bioinformatics 5: 59.

Krzywinski M, Schein J, Birol İ, Connors J, Gascoyne R, Horsman D, Jones SJ, Marra MA. 2009. Circos: An information aesthetic for comparative genomics. Genome Research 19: 1639–1645.

Lengyel S, Gove AD, Latimer AM, Majer JD, Dunn RR. 2010. Convergent evolution of seed dispersal by ants, and phylogeny and biogeography in flowering plants: a global survey. Perspectives in Plant Ecology 12: 43–55.

Mark S, Olesen JM. 1996. Importance of elaiosome size to removal of ant-dispersed seeds. Oecologia 107: 95–101.

Mendes FK, Vanderpool D, Fulton B, Hahn MW. 2021. CAFE 5 models variation in evolutionary rates among gene families. Bioinformatics 36: 5516–5518.

Miller CN, Whitehead SR, Kwit C. 2020. Effects of seed morphology and elaiosome chemical composition on attractiveness of five *Trillium* species to seed-dispersing ants. Ecology and Evolution 10: 2860–2873.

Murrell B, Wertheim JO, Moola S, Weighill T, Scheffler K, Pond SLK. 2012. Detecting Individual Sites Subject to Episodic Diversifying Selection. PLOS Genetics 8: e1002764.

Ou S, Su W, Liao Y, Chougule K, Agda JRA, Hellinga AJ, Lugo CSB, Elliott TA, Ware D, Peterson T, et al. 2019. Benchmarking transposable element annotation methods for creation of a streamlined, comprehensive pipeline. Genome Biology 20: 275.

Pertea M, et al. 2016. Transcript-level expression analysis of RNA-seq experiments with HISAT, StringTie and Ballgown. Nature Protocols 11: 1650–1667.

Pfeiffer M, Huttenlocher H, Ayasse M. 2010. Myrmecochorous plants use chemical mimicry to cheat seed-dispersing ants. Functional Ecology 24: 545–555.

Price MN, Dehal PS, Arkin AP. 2010. FastTree 2 – Approximately Maximum-Likelihood Trees for Large Alignments. PLOS ONE 5: e9490.

Qin RM, Wen P, Corlett RT, Zhang Y, Wang G, Chen J. 2022. Plant-defense mimicry facilitates rapid dispersal of short-lived seeds by hornets. Current Biology 32: 3429–3435.

Rambaut A, Drummond AJ, Xie D, Baele G, Suchard MA. 2018. Posterior Summarization in Bayesian Phylogenetics Using Tracer 1.7. Systematic Biology 67: 901–904.

Ramírez-Barahona S, Sauquet H, Magallón S. 2020. The delayed and geographically heterogeneous diversification of flowering plant families. Nature Ecology & Evolution 4: 1232–1238.

Schlüter PM, Xu S, Gagliardini V, Whittle E, Shanklin J, Grossniklaus U, Schiestl FP. 2011. Stearoyl-acyl carrier protein desaturases are associated with floral isolation in sexually deceptive orchids. Proceedings of the National Academy of Sciences of the U.S.A. 108, 5696–5701.

Schupp EW, Jordano P, Gómez JM. 2010. Seed dispersal effectiveness revisited: a conceptual review. New Phytologist 188: 333–353.

Sedeek KE, Whittle E, Guthörl D, Grossniklaus U, Shanklin J, Schlüter PM. 2016. Amino acid change in an orchid desaturase enables mimicry of the pollinator’s sex pheromone. Current Biology 26: 1505–1511.

Skidmore BA, Heithaus ER. 1988. Lipid cues for seed-carrying by ants in *Hepatica americana*. Journal of Chemical Ecology 14: 2185–2196.

Simão FA, Waterhouse RM, Ioannidis P, et al. 2015. BUSCO: assessing genome assembly and annotation completeness with single-copy orthologs. Bioinformatics 31: 3210–3212.

Smith MD, Wertheim JO, Weaver S, Murrell B, Scheffler K, Kosakovsky Pond SL. 2015. Less Is More: An Adaptive Branch-Site Random Effects Model for Efficient Detection of Episodic Diversifying Selection. Molecular Biology and Evolution 32: 1342–1353.

Smith SA, Brown JW, Walker JF. 2018. So many genes, so little time: A practical approach to divergence-time estimation in the genomic era. PLOS ONE 13: e0197433.

Smith A, Hubley R, Green P. 2013. RepeatMasker Open-4.0. URL: http://www.repeatmasker.org

Stamatakis A. 2014. RAxML version 8: a tool for phylogenetic analysis and post-analysis of large phylogenies. Bioinformatics 30: 1312–1313.

Stanke M, Keller O, Gunduz I, Hayes A, Waack S, Morgenstern B. 2006. AUGUSTUS: ab initio prediction of alternative transcripts. Nucleic Acids Research 34: W435–W439.

Tang H, Krishnakumar V, Zeng X, Xu Z, Taranto A, Lomas JS, Zhang Y, Huang Y, Wang Y, Yim WC, et al. 2024. JCVI: A versatile toolkit for comparative genomics analysis. iMeta 3: e211.

Tiffney BH. 2004. Vertebrate dispersal of seed plants through time. Annual Review of Ecology Evolution and Systematics 35: 1–29.

Turchetto-Zolet AC, Christoff AP, Kulcheski FR, Loss-Morais G, Margis R, Margis-Pinheiro M. 2016. Diversity and evolution of plant diacylglycerol acyltransferase (DGATs) unveiled by phylogenetic, gene structure and expression analyses. Genetics and Molecular Biology 39: 524–538.

Van Dongen S. 2008. Graph Clustering Via a Discrete Uncoupling Process. SIAM Journal on Matrix Analysis and Applications 30: 121–141.

Voelker TA, Worrell AC, Anderson L, Bleibaum J, Fan C, Hawkins DJ, Radke SE, Davies HM. 1992. Fatty acid biosynthesis redirected to medium chains in transgenic oilseed plants. Science 257: 72–74.

Walker BJ, et al. 2014. Pilon: an integrated tool for comprehensive microbial variant detection and genome assembly improvement. PLoS One 9: e112963.

Willson MF, Traveset A. 2000. The ecology of seed dispersal. In: Seeds: the ecology of regeneration in plant communities 2: 85–110.

Wong TKF, Ly-Trong N, Ren H, Baños H, Roger AJ, Susko E, Bielow C, Maio ND, Goldman N, Hahn MW, et al. 2025. IQ-TREE 3: Phylogenomic Inference Software using Complex Evolutionary Models.

Yang Z, Rannala B. 2006. Bayesian Estimation of Species Divergence Times Under a Molecular Clock Using Multiple Fossil Calibrations with Soft Bounds. Molecular Biology and Evolution 23: 212–226.

Yang Y, Smith SA. 2014. Orthology Inference in Nonmodel Organisms Using Transcriptomes and Low-Coverage Genomes: Improving Accuracy and Matrix Occupancy for Phylogenomics. Molecular Biology and Evolution 31: 3081–3092.

Yang T, Niu Q, Dai H, Tian X, Ma J, Pritchard HW, Lin L, Yang X. 2024. The transcription factor MYB1 activates DGAT2 transcription to promote triacylglycerol accumulation in sacha inchi (*Plukenetia volubilis* L.) leaves under heat stress. Plant Physiology and Biochemistry 208: 108517.

Yu YL, Chomicki G, Chen, G. 2025. Seed dispersal by kleptoparasitic flies. Current Biology, 35: R700–R701.

Zhang R-G, Li G-Y, Wang X-L, Dainat J, Wang Z-X, Ou S, Ma Y. 2022. TEsorter: An accurate and fast method to classify LTR-retrotransposons in plant genomes. Horticulture Research 9: uhac017.

Zhang Y, Li Y, Han B, Liu A, Xu W. 2022. Integrated lipidomic and transcriptomic analysis reveals triacylglycerol accumulation in castor bean seedlings under heat stress. Industrial Crops & Products 180: 114702.

Zhang C, Nielsen R, Mirarab S. 2025. ASTER: A Package for Large-Scale Phylogenomic Reconstructions. Molecular Biology and Evolution 42: msaf172.

Doležel J, Greilhuber J, Suda J. 2007. Estimation of nuclear DNA content in plants using flow cytometry. Nature Protocols 2: 2233–2244.

Durand NC, Shamim MS, Machol I, Rao SS, Huntley MH, Lander ES, Aiden EL. 2016. Juicer Provides a One-Click System for Analyzing Loop-Resolution Hi-C Experiments. Cell Systems 3: 95–98.

Dudchenko O, Batra SS, Omer AD, Nyquist SK, Hoeger M, Durand NC, Shamim MS, Machol I, Lander ES, Aiden AP, Aiden EL. 2017. De novo assembly of the Aedes aegypti genome using Hi-C yields chromosome-length scaffolds. Science 356: 92–95.

Ranallo-Benavidez TR, Jaron KS, Schatz MC. 2020. GenomeScope 2.0 and Smudgeplot for reference-free profiling of polyploid genomes. Nature Communications 11:1432.

Kokot M, Dlugosz M, Deorowicz S. 2017. KMC 3: counting and manipulating k-mer statistics. Bioinformatics 33: 2759–2761.

Xu M, Guo L, Gu S, Wang O, Zhang R, Peters BA, Fan G, Liu X, Xu X, Deng L, Zhang Y. 2020. TGS-GapCloser: A fast and accurate gap closer for large genomes with low coverage of error-prone long reads. Gigascience 9: giaa094.

Hu J, Wang Z, Liang F, Liu SL, Ye K, Wang DP. 2024. NextPolish2: A Repeat-aware Polishing Tool for Genomes Assembled Using HiFi Long Reads. Genomics Proteomics Bioinformatics 22: qzad009.

