## Supplementary Materials and Methods and Figs S1-S6 for "*Stemona* genomes illuminate fatty acid partitioning between seeds and elaiosomes mediating wasp dispersal"

### Genome annotation and comparative genomics

For both genomes, we first annotated repetitive elements using EDTA v2.22 (--sensitive 1 --anno 1), to generate a *de novo* library of repetitive elements per genome. RepeatMasker v4.18 was subsequently applied to softmask repetitive sequences prior to protein-coding gene annotation. To annotate protein-coding genes we combined RNA-seq (see *Transcriptome sequencing and analyses*, below) and homologous protein evidence from published genomes (Tables S1-2). Short read RNA-seq libraries were combined and assembled with Trinity v2.15.2 (Grabherr *et al.*, 2011) and were further aligned to the assemblies with HISAT2 v2.2.1 (Pertea *et al.*, 2016) and assembled into transcripts with StringTie v2.1.4 (Pertea *et al.*, 2016). Transcript assemblies were combined using PASA v2.5.3 (Haas *et al.*, 2008). Full length ORFs were identified with TransDecoder v5.7.0 (https://github.com/TransDecoder/TransDecoder) and high-quality gene models identified by BLAST searches to the homologous protein databases were used to train SNAP (Korf, 2004) and AUGUSTUS v3.5.0 (Stanke *et al.*, 2006) with 5 rounds of training. Additionally, MAKER2 v3.01.03 (Holt & Yandell, 2011) was used to annotate the assembly leveraging *ab initio* SNAP and AUGUSTUS predictions, PASA transcript assemblies and homologous proteins. Finally, EVidenceModeler v2.1.0 (Haas *et al.*, 2008) was used to integrate MAKER annotations and PASA transcript assemblies, masking any additional protein-coding repetitive elements identified by TEsorter v1.4.7 (Zhang *et al.*, 2022). EVidenceModeler annotations were updated with PASA to add UTRs and alternative splicing, for two rounds. This resulted in some loci being erroneously merged, and these were identified and manually curated. Incomplete gene models and ORFs shorter than 50aa were removed and overlapping genes on the same strand were filtered. The completeness of the final annotation was assessed with BUSCO (-m prot) with Embryophyta odb10.

We compared macrosynteny between our two newly assembled species using the MCScan algorithm as implemented in JCVI (Tang *et al*., 2024). Simplified macrosynteny blocks were combined with GC content, repeat count, and gene count in 100 Kb windows and plotted using circos v0.69-10 (Krzywinski *et al*., 2009).

We assembled a phylogenomic dataset for Pandanales by combining annotation reference genomes with additional published genome and transcriptomes assemblies (Tables S1-2). Where necessary, RNA-seq data was combined and assembled with Trinity and ORFs were predicted with TransDecoder. For *Xerophyta viscosa* (Velloziaceae) and *Pandanus amaryllifolius* (Pandanaceae), no annotations were available, and these assemblies were first annotated with BRAKER3 (Gabriel *et al*., 2024) using published RNA-seq data and the same homologous protein evidence as above (Tables S1-2). Genome annotations were filtered to retain a single primary transcript per locus, and transcriptome ORFs were clustered with CD-HIT v4.8.1 (Fu et al., 2012) (-c 0.995). Due to the inclusion of transcriptome data, we adopted the phylogenomic pipeline from Yang & Smith (2014), using scripts from https://bitbucket.org/yanglab/phylogenomic_dataset_construction. Peptide sequences were subjected to all-by-all similarity searches using DIAMOND v2.1.10 (Buchfink *et al*., 2021), preserving hits with at least 50% query coverage. Sequences were clustered using MCL v22.282 (van Dongen, 2008) (-tf 'gq(5)' -I 1.5) and clusters with at least 10 taxa represented were retained. Clusters were aligned with MAFFT v7.505 (Katoh & Standley, 2013) (--auto), columns with less than 10% sequence occupancy were removed with pxclsq from phyx v1.3 (Brown *et al*., 2017), and trees were inferred with FastTree v2.1.11 (Price *et al*., 2010) (-wag). Tips longer than 1.0 substitution per site or longer than 0.5 and > 10x their sister were removed. Clusters of monophyletic sequences from transcriptomes were masked to the single longest sequence in the cleaned alignment. Finally, clusters subtended by branches longer than 1.0 substitution per site were pruned and retained if they contained at least 10 sequences. All steps were then repeated two further times. Finally, peptide sequences were aligned with MAFFT (--maxiterate 1000 --genafpair), coding sequences were aligned to proteins with pxaa2cdn from phyx, low occupancy columns were removed as above and trees were inferred with IQ-TREE v3.0.1 (Wong *et al*., 2025) (-m GTR+F+G). Monophyletic clusters of sequences from a single taxon were collapsed to the one on the shortest branch, and one-to-one and Minimum Inclusion orthologues were pruned from these trees. Gene trees from orthologues with full taxon occupancy were used to infer a species tree with ASTRAL-IV v1.23.4.6, implemented in ASTER (Zhang *et al*., 2025). We used the gene shopping approach implemented in SortaDate (Smith *et al*., 2018) to winnow this dataset for divergence time inference, selecting 10 orthologues that were perfectly concordant with the species tree and ranking based on root-to-tip variance and then tree length. Coding sequence alignments for these genes were concatenated and used to infer divergence times on the species tree with mcmctree in PAML v4.10.9 (Yang & Rannala, 2006) using the approximate likelihood approach (dos Reis & Yang 2011). We used a series of secondary calibrations based on the conservative complete analysis of Ramirez-Barahona *et al*. (2020). We used the root age constraint and the average molecular root height to calibrate the locus rate prior with shape 2 and scale 5.23 and set a gamma prior on the rate variance with shape 1 and scale 10. We used a Birth-Death tree prior with birth = death = 1.0 and a sampling fraction of 0.1, and the multiplicative construction of the effective prior. We ran MCMC for 2 million iterations, sampling every thousand, and burned in the first 100,000 iterations. We used Tracer v1.7 (Rambaut *et al*., 2018) to calculate Effective Sample Sizes (ESS) for numerical parameters and ensured that all ESS > 200.

We took gene counts per taxon from our final homologue groups prior to pruning orthologues and used them to infer gene family expansions and contractions under a

Birth-Death model with CAFE5 v1.1 (Mendes *et al.*, 2020). To ensure convergence, we ranked homologues by count variance and excluded the 11 most variable. We inferred an error model to account for the fact that our dataset includes transcriptome data where apparent loss can be due to lack of expression.

To search for functional diversification associated with transitions to myrmecochory or vespicochory, we fit codon models to test for diversifying selection along the gene trees of our candidate functional genes (see below). Final homologue clusters containing each characterised *S. tuberosa* gene were collected, coding sequences were aligned, columns with less than 80% occupancy were removed, and gene trees were inferred as above. To reduce computational and numerical complexity, we collapsed monophyletic clusters representing species-specific duplications from species outside our focal pair. We used ABSREL implemented in HyPhy v2.5.72 (Smith *et al.*, 2015), accounting for synonymous rate variation. We specified foreground branches to test as: i) the branch subtending the MRCA of *Stemona* and *Croomia*; ii) the branch subtending *Stemona*; and iii) the branch subtending *S. tuberosa*. If lineage-specific paralogues were present in the tree, labelling was applied the same way to each one, except that the branch subtending an implied duplication node was also labelled as foreground. Some truncated annotations created regions of poor alignment due to lack of homology, and these were masked in foreground sequences by replacing the relevant bases with strings of Ns. We additionally removed any columns where a majority of foreground sequences were missing data. We fit the models and searched for branches with Holm-Bonferroni adjusted p-values $< 0.05$. To infer per-site evidence of diversifying selection along branches with significant results from ABSREL, we applied MEME (Murrell *et al.*, 2012), specifying as foreground the significant branches and applying a p-value cutoff of 0.05.

**References:**

**Brown JW, Walker JF, Smith SA. 2017.** Phyx: phylogenetic tools for unix.

*Bioinformatics* 33: 1886–1888.

**Buchfink B, Reuter K, Drost H-G. 2021.** Sensitive protein alignments at tree-of-life scale using DIAMOND. *Nature Methods* 18: 366–368.

**dos Reis M, Yang Z. 2011.** Approximate Likelihood Calculation on a Phylogeny for Bayesian Estimation of Divergence Times. *Molecular Biology and Evolution* 28: 2161–2172.

**Fu L, Niu B, Zhu Z, Wu S, Li W. 2012.** CD-HIT: accelerated for clustering the next-generation sequencing data. *Bioinformatics* 28: 3150–3152.

**Gabriel L, Brůna T, Hoff KJ, Ebel M, Lomsadze A, Borodovsky M, Stanke M. 2024.** BRAKER3: Fully automated genome annotation using RNA-seq and protein evidence with GeneMark-ETP, AUGUSTUS, and TSEBRA. Genome Research 34: 769–777.

**Grabherr MG, Haas BJ, Yassour M, Levin JZ, Thompson DA, Amit I, Adiconis X, Fan L, Raychowdhury R, Zeng Q *et al.* 2011.** Full-length transcriptome assembly from RNA-Seq data without a reference genome. *Nature Biotechnology* 29: 644–652.

**Haas BJ, Salzberg SL, Zhu W, Pertea M, Allen JE, Orvis J, White O, Buell CR, Wortman JR. 2008.** Automated eukaryotic gene structure annotation using EVidenceModeler and the Program to Assemble Spliced Alignments. *Genome Biology* 9: R7.

**Holt C, Yandell M. 2011.** MAKER2: an annotation pipeline and genome-database management tool for second-generation genome projects. *BMC Bioinformatics* 12: 491.

**Katoh K, Standley DM. 2013.** MAFFT Multiple Sequence Alignment Software Version 7: Improvements in Performance and Usability. *Molecular Biology and Evolution* 30: 772–780.

**Korf I. 2004.** Gene finding in novel genomes. *BMC Bioinformatics* 5: 59.

**Krzywinski M, Schein J, Birol İ, Connors J, Gascoyne R, Horsman D, Jones SJ, Marra MA. 2009.** Circos: An information aesthetic for comparative genomics. *Genome Research* 19: 1639–1645.

**Mendes FK, Vanderpool D, Fulton B, Hahn MW. 2021.** CAFE 5 models variation in evolutionary rates among gene families. *Bioinformatics* 36: 5516–5518.

**Murrell B, Wertheim JO, Moola S, Weighill T, Scheffler K, Pond SLK. 2012.** Detecting Individual Sites Subject to Episodic Diversifying Selection. *PLOS Genetics* 8: e1002764.

**Pertea M, Kim D, Pertea GM, Leek JT, Salzberg SL. 2016.** Transcript-level expression analysis of RNA-seq experiments with HISAT, StringTie and Ballgown. *Nature Protocols* 11: 1650–1667.

**Price MN, Dehal PS, Arkin AP. 2010.** FastTree 2 – Approximately Maximum-

Likelihood Trees for Large Alignments. *PLOS ONE* 5: e9490.

**Rambaut A, Drummond AJ, Xie D, Baele G, Suchard MA. 2018.** Posterior Summarization in Bayesian Phylogenetics Using Tracer 1.7. *Systematic Biology* 67: 901–904.

**Ramírez-Barahona S, Sauquet H, Magallón S. 2020.** The delayed and geographically heterogeneous diversification of flowering plant families. *Nature Ecology & Evolution* 4: 1232–1238.

**Smith MD, Wertheim JO, Weaver S, Murrell B, Scheffler K, Kosakovsky Pond SL. 2015.** Less Is More: An Adaptive Branch-Site Random Effects Model for Efficient Detection of Episodic Diversifying Selection. *Molecular Biology and Evolution* 32: 1342–1353.

**Smith SA, Brown JW, Walker JF. 2018.** So many genes, so little time: A practical approach to divergence-time estimation in the genomic era. *PLOS ONE* 13: e0197433.

**Stanke M, Keller O, Gunduz I, Hayes A, Waack S, Morgenstern B. 2006.** AUGUSTUS: ab initio prediction of alternative transcripts. *Nucleic Acids Research* 34: W435–W439.

**Tang H, Krishnakumar V, Zeng X, Xu Z, Taranto A, Lomas JS, Zhang Y, Huang Y, Wang Y, Yim WC et al. 2024.** JCVI: A versatile toolkit for comparative genomics analysis. *iMeta* 3: e211.

**Van Dongen S. 2008.** Graph Clustering Via a Discrete Uncoupling Process. *SIAM Journal on Matrix Analysis and Applications* 30: 121–141.

**Wong TKF, Ly-Trong N, Ren H, Baños H, Roger AJ, Susko E, Bielow C, Maio ND, Goldman N, Hahn MW et al. 2025.** IQ-TREE 3: Phylogenomic Inference Software using Complex Evolutionary Models. *EcoEvoRxiv.* https://doi.org/10.32942/X2P62N

**Yang Y, Smith SA. 2014.** Orthology Inference in Nonmodel Organisms Using Transcriptomes and Low-Coverage Genomes: Improving Accuracy and Matrix Occupancy for Phylogenomics. *Molecular Biology and Evolution* 31: 3081–3092.

**Yang Z, Rannala B. 2006.** Bayesian Estimation of Species Divergence Times Under a Molecular Clock Using Multiple Fossil Calibrations with Soft Bounds. *Molecular Biology and Evolution* 23: 212–226.

**Zhang C, Nielsen R, Mirarab S. 2025.** ASTER: A Package for Large-Scale Phylogenomic Reconstructions. *Molecular Biology and Evolution* 42: 1-4.

**Zhang R-G, Li G-Y, Wang X-L, Dainat J, Wang Z-X, Ou S, Ma Y. 2022**. TEsorter: An accurate and fast method to classify LTR-retrotransposons in plant genomes. *Horticulture Research* 9: uhac017.

**Supplementary figures:**

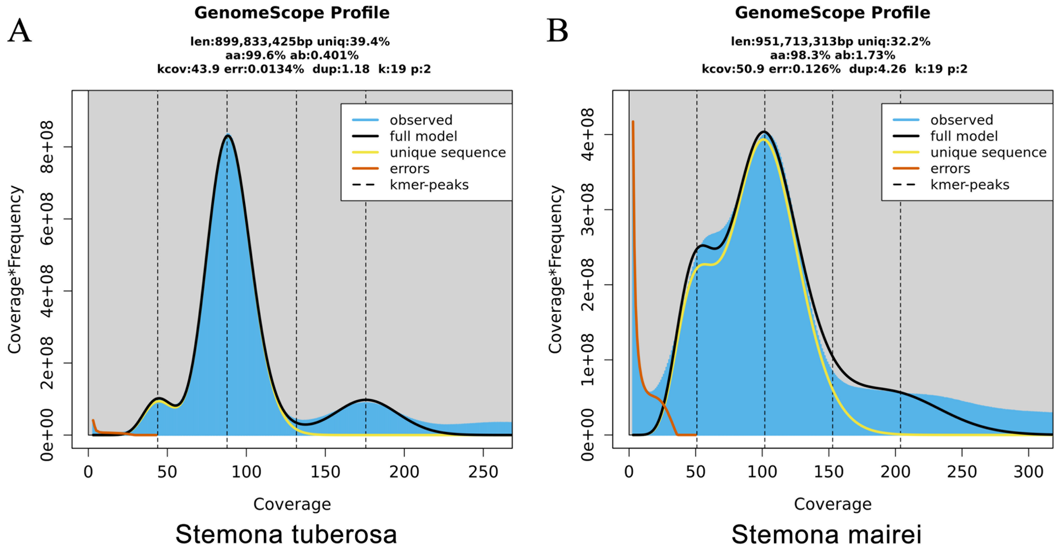


**Figure S1.** GenomeScope 2.0 k-mer frequency profiles for (A) *Stemona tuberosa* and (B) *S. mairei*, based on Illumina short-read data with k = 19 and ploidy = 2. The x-axis shows k-mer coverage depth and the y-axis shows coverage x frequency. The observed k-mer spectrum (blue), fitted model (black), unique sequence component (yellow), and error component (orange) are shown; dashed vertical lines mark the heterozygous and homozygous k-mer coverage peaks. Estimated genome sizes were 899.8 Mb for S. tuberosa (heterozygosity 0.401%, error rate 0.013%, duplication rate 1.18) and 951.7 Mb for *S. mairei* (heterozygosity 1.73%, error rate 0.126%, duplication rate 4.26). The more prominent heterozygous peak and higher duplication rate in *S. mairei* are consistent with its larger genome and greater repeat content.

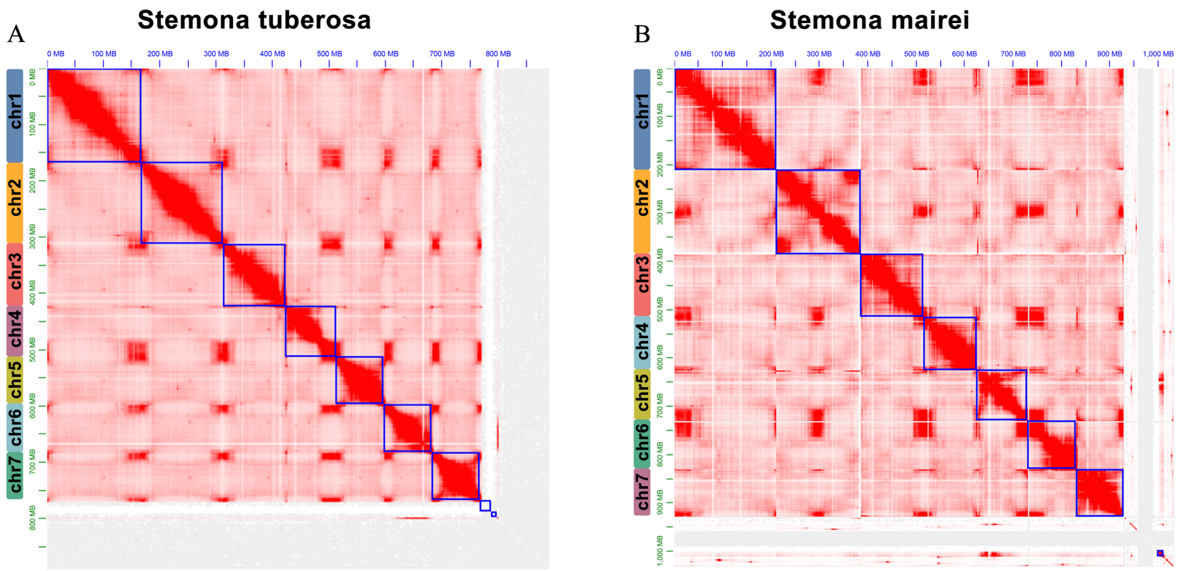


**Figure S2.** Hi-C map (High-throughput chromosome conformation capture) of the *Stemona tuberosa* (a) and *S. mairei* (b) genomes showing strong interactive signal within intra-chromosome and at diagonal regions. The color bar indicates the interactive signal.

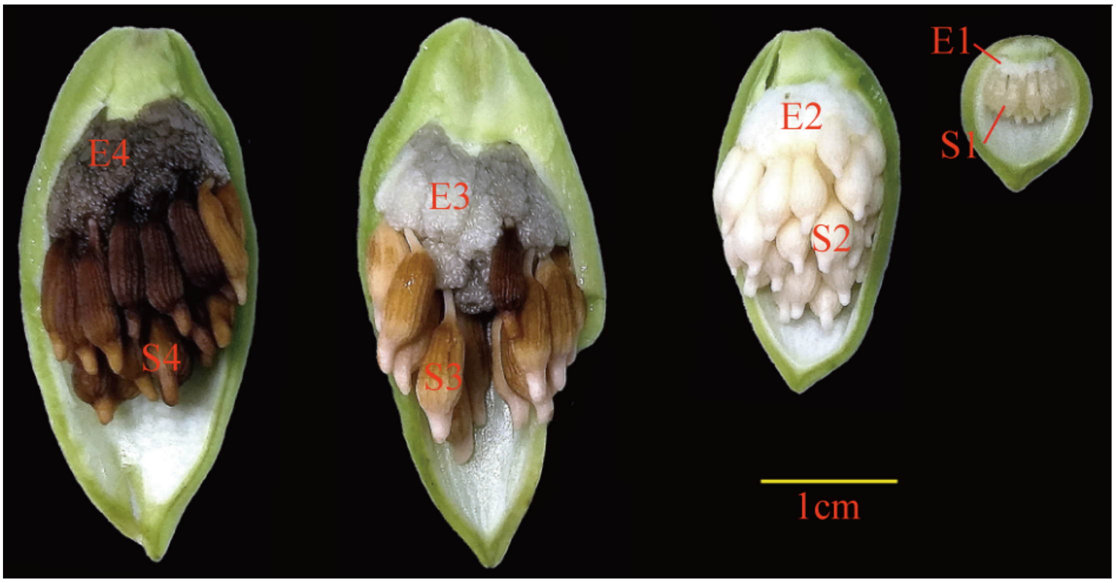


**Figure S3.** The shape, size and colors of seed (S1-S4) and elaiosome (E1-E4) during fruit development and ripening at different stages. Bar, 1 cm.

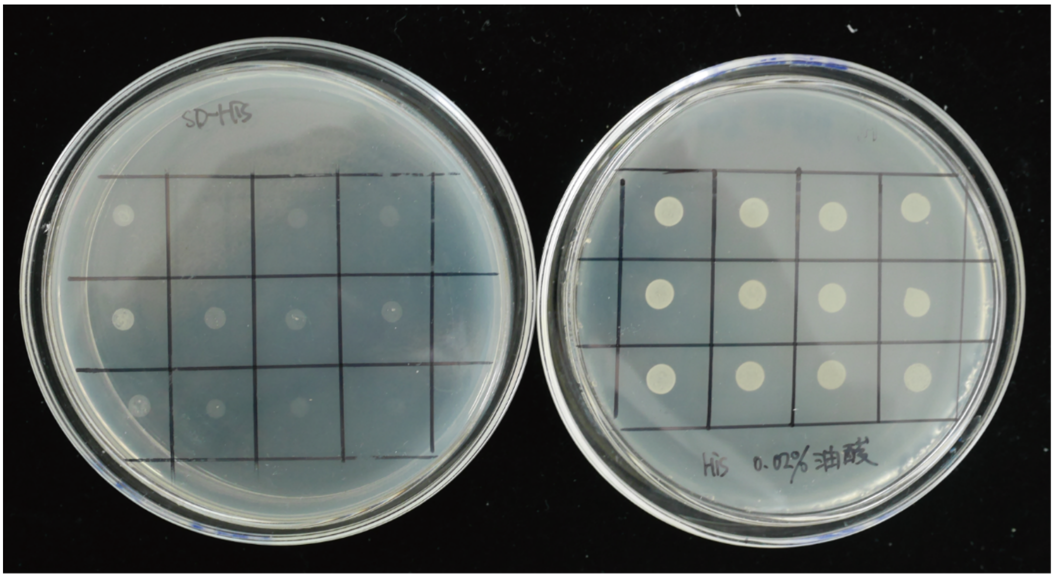


**Figure S4.** Live/dead spot assay of yeast BY4741 mutant strain *Δole1* (*MATa ole1Δ::HIS3 leu2Δ0 met15Δ0 ura3Δ0*) on synthetic complete selection medium (-His) with (right panel) or without (left panel) oleic acid supplement.

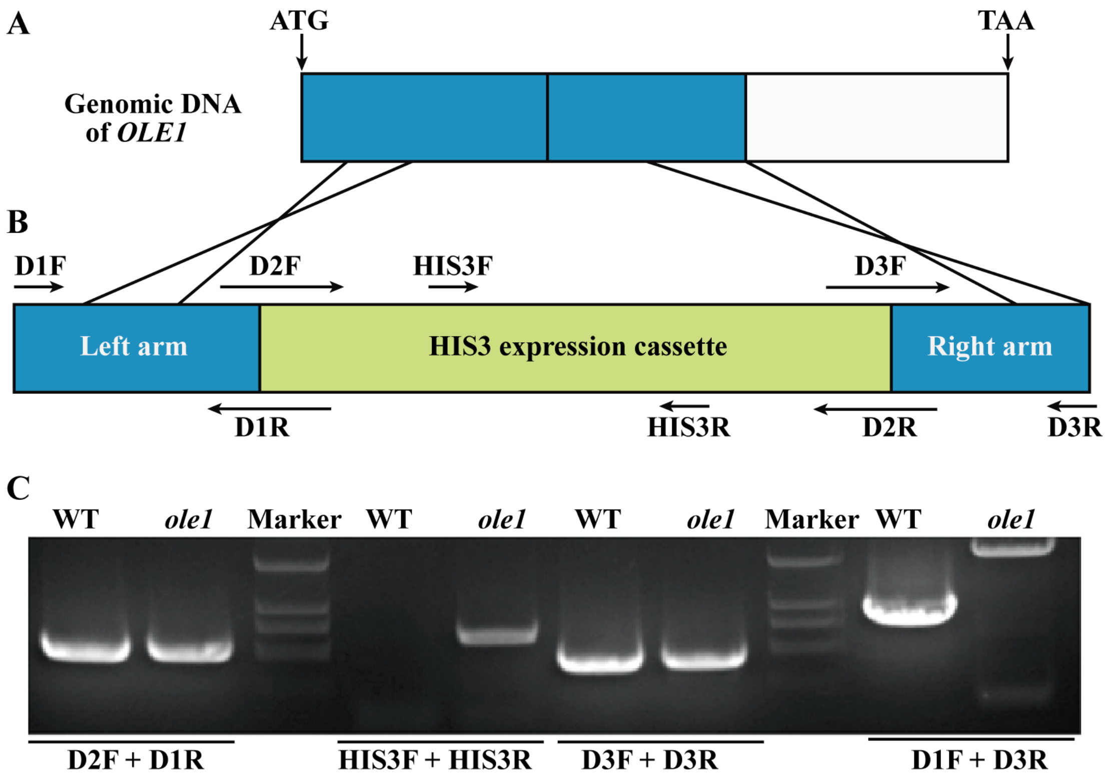


**Figure S5**. Construction and molecular identification of *OLE1* mutant (*Δole1*) in yeast strain BY4741.

(A). Schematic representation of *OLE1* genomic DNA from ATG to TAA. The black line in the blue box shows the region that will be disrupted.

(B). Schematic representation of *OLE1* deletion cassette. The left and right arm (light blue box) are homologous regions of *OLE1* genomic DNA as shown in A. The light green box represents the HIS3 expression cassette. Specific primers used for construction of *OLE1* deletion cassette and molecular identification for yeast strain BY474141 and *Δole1* are indicated by the arrows.

(C). Polymerase chain reaction (PCR)-based identification of *Δole1* mutant line with specific primers.

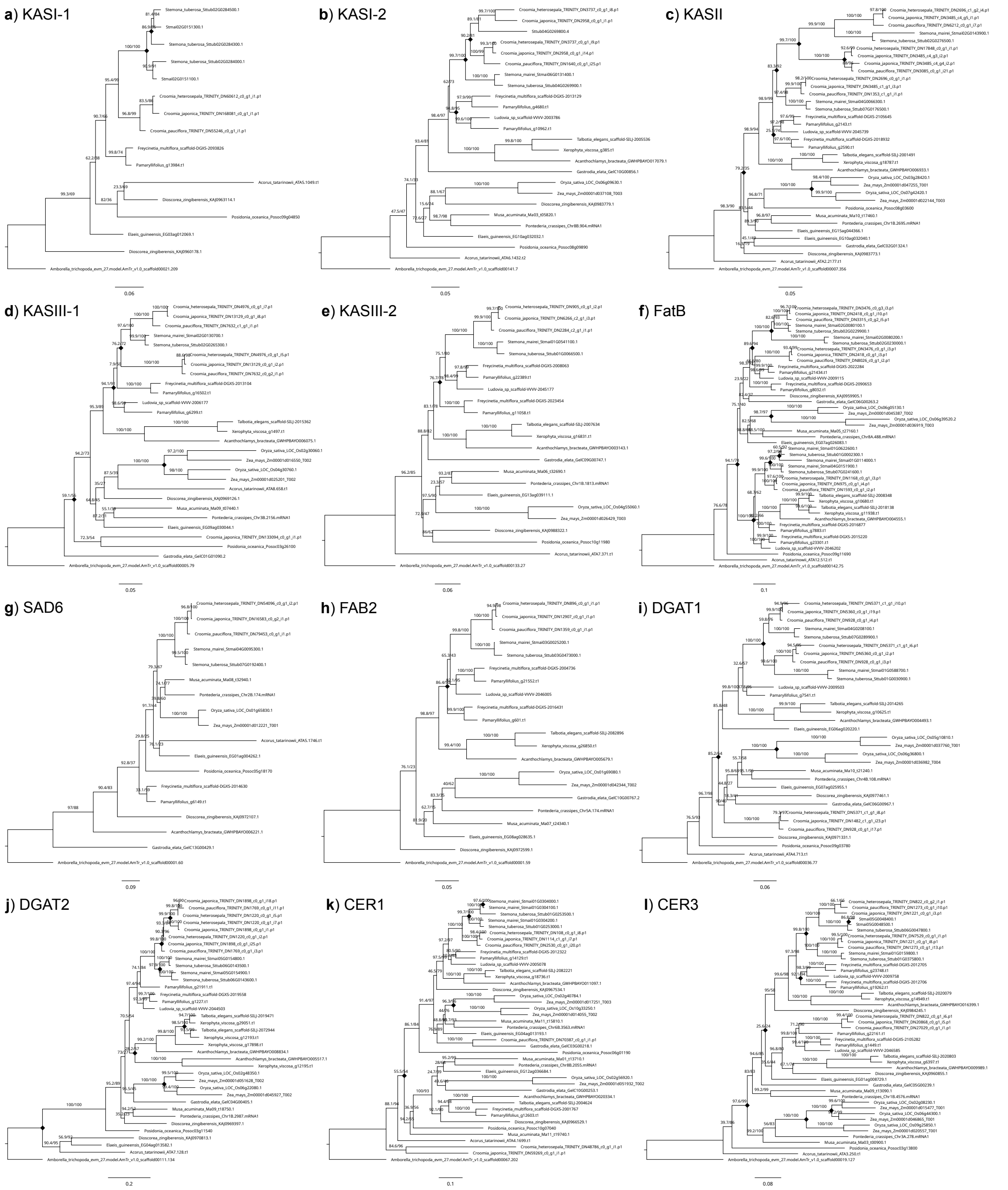


**Figure S6.** Gene trees of the main fatty acid synthesis gene families involved in the differentiation of elaiosome and seed lipids. a) KASI-1, b) KASI-2, c) KASII, d) KASIII-1, e) KASIII-2, f) FatB, g) SAD6, h) FAB2, i) SGAT1, j) DGAT2, k) CER1, i) CER3. Diamonds at nodes indicate inferred gene duplications based on species overlaps. Scale bars give the expected number of nucleotide substitutions per site.
